# Chemical screens in aging-relevant human motor neurons identify MAP4Ks as therapeutic targets for amyotrophic lateral sclerosis

**DOI:** 10.1101/2023.04.24.538014

**Authors:** Meng-Lu Liu, Shuaipeng Ma, Wenjiao Tai, Xiaoling Zhong, Haoqi Ni, Yuhua Zou, Jingcheng Wang, Chun-Li Zhang

## Abstract

Effective therapeutics is much needed for amyotrophic lateral sclerosis (ALS), an adult-onset neurodegenerative disease mainly affecting motor neurons. By screening chemical compounds in human patient-derived and aging-relevant motor neurons, we identify a neuroprotective compound and show that MAP4Ks may serve as therapeutic targets for treating ALS. The lead compound broadly improves survival and function of motor neurons directly converted from human ALS patients. Mechanistically, it works as an inhibitor of MAP4Ks, regulates the MAP4Ks-HDAC6-TUBA4A-RANGAP1 pathway, and normalizes subcellular distribution of RANGAP1 and TDP-43. Finally, in an ALS mouse model we show that inhibiting MAP4Ks preserves motor neurons and significantly extends animal lifespan.

## INTRODUCTION

Amyotrophic lateral sclerosis (ALS) is an adult-onset neurodegenerative disease, featuring progressive death of motor neurons (MNs) and loss of the ability to walk, talk, swallow and breathe. Life expectancy for ALS patients is about 2-5 years. So far, there is no effective treatment, begging for discovery of potential therapeutic targets.

Because of the unique cellular pathology of ALS, aging-relevant human motor neurons will be most appropriate for therapeutic identification. However, it is impossible to isolate such live neurons directly from human ALS patients. As an alternative, one emerging strategy is to directly convert non-neuronal somatic cells into neurons. This is largely accomplished via manipulation of key fate-determining factors (1). Through such a strategy, human skin fibroblasts can be directly converted into diverse subtypes of neurons in culture, such as cortical neurons (2, 3), dopaminergic neurons (4, 5), medium spiny neurons (6), cholinergic neurons (7, 8), motor neurons (9, 10), and serotonergic neurons (11, 12). One distinct feature of these directly converted human neurons is that they maintain aging-associated features, in contrasts to those that are derived from induced pluripotent stem cells (iPSCs) which are age-reset to an embryonic stage (13–18). Stem cell-derived neurons are transcriptionally immature and do not exhibit hallmarks of neurodegenerative diseases (19). This is likely due to that the aging process leads to accumulation of stresses that are associated with protein aggregations and dysregulation of signaling pathways, nucleocytoplasmic transports and epigenetics. Comparing to non-neuronal cells or stem cell-derived neurons, aging-relevant and disease-associated human neurons may be better substrates for therapeutic identifications.

Previously, we showed that the small molecule forskolin and the transcription factor NEUROG2 could convert human skin fibroblasts into cholinergic neurons (8). They could be further specified to become motor neurons by two additional transcription factors, ISL1 and LHX3, during the conversion process (10). These human induced motor neurons (hiMNs) exhibited electrophysiological features of mature neurons and formed functional neuromuscular junctions (NMJs) in a co-culture system (10). Although fibroblasts from both healthy and ALS patients could be equally converted, ALS-hiMNs showed pathology including smaller soma, simpler neurites, fewer NMJs when co-cultured with skeletal muscles, and poor survival (10). Importantly, these hiMNs could be used in cell-based assays, evidenced by our prior disease modeling studies (10, 20, 21) and candidate-based identification of kenpaullone as a small molecule capable of promoting survival and function of ALS-hiMNs (10).

In this study, we further optimized hiMN-based survival assays and conducted high-throughput screens of a chemical library of bioactive small molecules using aging-relevant ALS-hiMNs for the first time. We then systematically studied the top hit chemical compound and revealed its target and the downstream signaling pathway. Validating our approach is that the target kinases, MAP4Ks, were previously implicated in neurodegeneration when studied in stem cell-derived neurons (19, 22–24). However, our study revealed a novel downstream signaling pathway, reflecting the key biological differences between embryonic and aging-relevant neurons. Furthermore, we examined the in vivo function of targeting this pathway in the mouse SOD1^G93A^ model of ALS and found that animal lifespan could be significantly extended after chemical treatment.

## RESULTS

### A compound screening platform with ALS-hiMNs

Using ALS1-hiMNs (ND29563, 37 years, designated as ALS1, Table S1), our prior pilot screen identified the GSK3β inhibitor kenpaullone (Ken) as a chemical compound with neuroprotective effect (10). We then modified this cell-based platform for screening more chemical compounds. Since reprogrammed cells are initially smaller in size and slower to attach to the plastic plates when compared to those non-reprogrammed fibroblasts, a replating step and passing through a 20-µm strainer enriched highly pure (> 85%, based on the co-expressed GFP) ALS1-hiMNs for cell viability assays (Fig. 1A). The ATP-based CellTiter-Glo luminescent assays exhibited good linearity over low (∼1,000 cells/well) to high (∼10,000 cells/well) seeding density of ALS1-hiMNs in 96-well plates. Each screen cycle could be finished three days post a single treatment of individual compound. Using Ken as a positive control, a Z-prime value ≥ 0.5 could be obtained for each assay.

**Figure 1.**
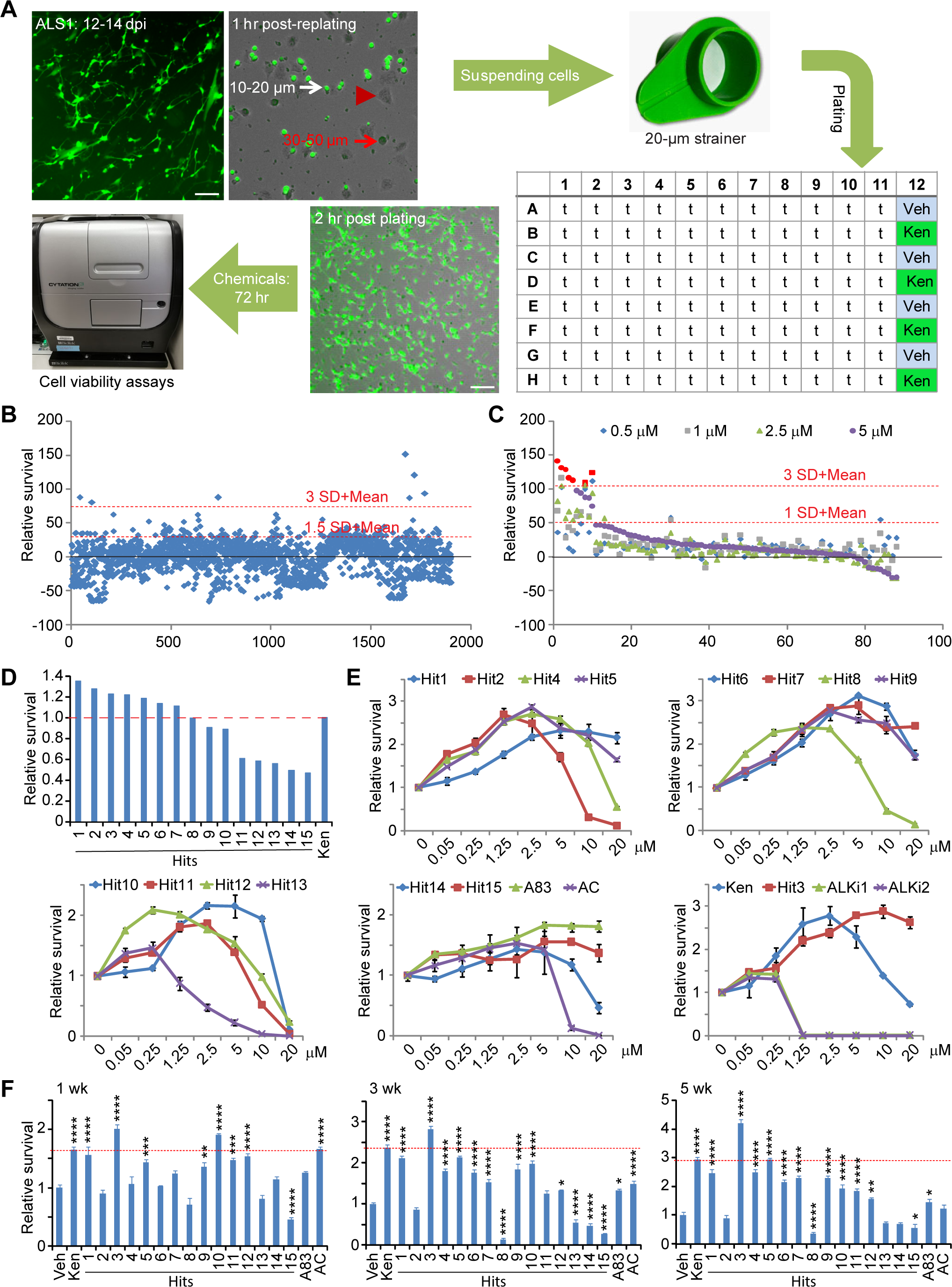
Screens for chemicals improving survival of ALS-hiMNs. A. A flowchart of the screening procedure. Highly pure (>85%) ALS1-hiMNs could be obtained after a replating procedure and passing through a cell strainer based on differential plate-attachment and cell size. The non-converted fibroblasts (indicated by an arrow or arrowhead) were larger and attached faster than the converted cells (indicated by a white arrow). An ATP-based assay was used to measure cell numbers. Veh (DMSO) and Ken (Kenpaullone) served as negative and positive controls, respectively. dpi, days post infection. Scale bar, 100 µm. B. Scatter plots of primary screens. Each point represents an individual compound assayed at 2.5 µM (final concentration). SD, standard deviation. C. Scatter plots of secondary screens. Top 65 hits from primary screens, together with 23 additional inhibitors targeting ALK, TGFβ, and GSK3, were examined at four different concentrations. D. Survival effect on ALS1-hiMNs by top 15 hits, including six inhibitors of GSK-3 (Hit1, 1-Azakenpaullone; Hit4, AZD1080; Hit5, AZD2858; Hit6, SB216763; Hit10, CHIR-99021; Hit13, BIO). Red line indicates the effect of the positive control Ken. E. Dose-response curves (mean ± SEM; n = 4 independent samples at each concentration). Top 15 hits from secondary screens, together with the indicated additional chemicals, were examined at 7 concentrations. F. Survival of ALS1-hiMNs when cocultured with astrocytes and treated with the indicated chemicals (mean ± SEM; n = 5 independent samples; *p < 0.05, **p < 0.01, ***p < 0.001, and ****p < 0.0001 when compared to the vehicle control, Student’s t-test). Red line indicates the effect of the positive control Ken. wk, week(s) on co-cultured astrocytes.

### Discovery of protective compounds for ALS-hiMNs

We then screened a library of about 2,000 compounds consisting of FDA-approved drugs and bioactive chemicals. The primary screens were conducted at 2.5 µM (final concentration) for each compound and the results were included in Table S2. We choose the top 65 chemicals (≥1.5 standard deviations above the mean of all tested compounds; Fig. 1B), together with 23 additional inhibitors targeting TGFβ (25–27), or GSK3 (28), for secondary screens at four concentrations (Fig. 1C). From secondary screens, we further assessed the top 15 chemicals (Fig. 1D; Table S3) in dose-response assays (Fig. 1E). Like Ken, most GSK3 inhibitors (Hit1, 4, 5, 6, and 9) as well as the ALK inhibitor K02288 (Hit3) could promote neuronal survival, although the optimal drug dosage varied for each chemical (Fig. 1E).

The above 15 chemicals were further examined through a tertiary screen, which was conducted with ALS1-hiMNs cocultured with primary astrocytes from wildtype mice. Cells were treated with these chemicals at their optimal dosage and were quantified by automated imaging of GFP+ hiMNs through a time-course (Fig. 1F). Unexpectedly, some chemicals no longer exhibited protections of ALS-hiMNs (Fig. 1F, 5 weeks on astrocytes), likely due to the neuronal supportive function of the co-cultured astrocytes that might mask the chemical’s minor effect. These chemicals were excluded from further analysis since neurons are normally supported by astrocytes under an in vivo condition. The targets of some of those chemicals continually showed neuronal protections are known to be associated with ALS, such as GSK3 (Ken, Hit1, 4, 5, 6, and 9) (28, 29) and TGFβ/BMP (Hit3) (26). Of note, several other TGFβ/BMP inhibitors, such as A83, AC (LDN-214117), LDN-193189 (ALKi1) and LDN-212854 (ALKi2) (30), were less effective (Fig. 1E, F). Overall, Hit3 showed the most protective effect on ALS1-hiMNs and was selected for subsequent analyses.

### Hit3 broadly protects ALS-hiMNs and improves their function

Using the neuron-glia coculture system, Hit3, along with Ken and two TGFβ/BMP inhibitors (A83 and ALKi), was examined on hiMNs derived from fibroblasts of healthy control patients (NL1) or patients with mutations on FUS (ALS1 and ALS2), SOD1 (ALS4), or C9ORF72 (ALS7 and ALS8) (Fig. 2A; Table S1). Hit3 treatments dosage-dependently improved survival of all these hiMNs (Fig. 2A), while the control TGFβ/BMP inhibitor either had no effect (A83) or was toxic (ALKi). Hit3 outperformed Ken, with the latter showing toxicity at higher concentrations (Fig. 2A, B).

**Figure 2.**
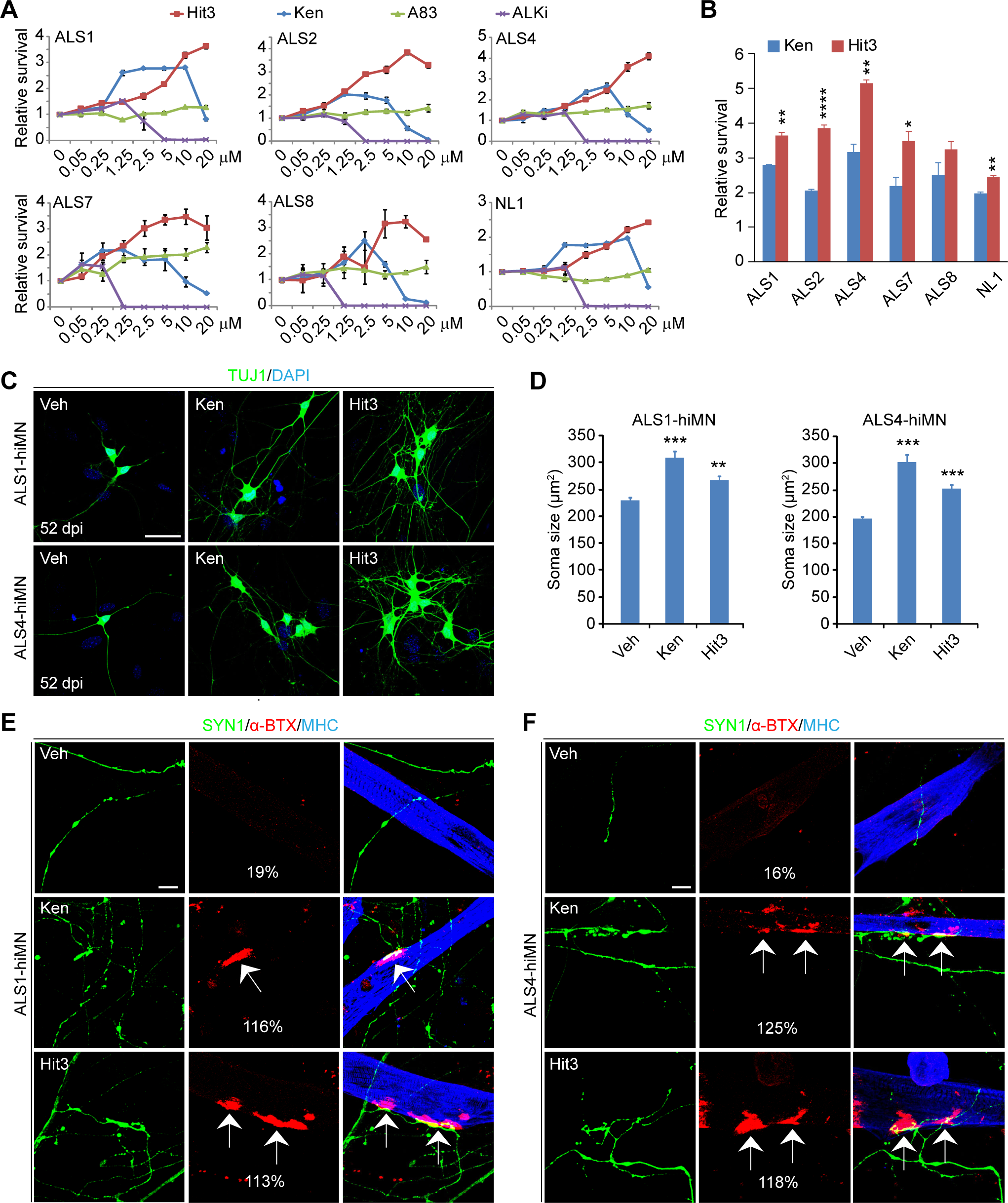
Hit3 improves survival and function of hiMNs from diverse human patients. A. Dose-dependent effects of the indicated chemicals on hiMNs from diverse ALS patients (mean ± SEM; n = 4 independent samples at each concentration). B. Hit3 outperforms the positive control Ken, in promoting survival of diverse ALS-hiMNs co-cultured with astrocytes (mean ± SEM; n = 4 independent samples at each concentration; *p < 0.05, **p < 0.01 and ****p < 0.0001 when compared to the Ken-treated samples, Student’s t-test). C. Hit3 and the positive control Ken increased neuronal soma size and complexity of ALS1-hiMNs and ALS4-hiMNs. Scale bar, 50 µm. D. Quantification of neuronal soma size. (mean ± SEM; n = 3 independent samples with ≥70 neurons analyzed for each group; **p < 0.01 and ***p < 0.001 when compared to the vehicle group, Student’s t-test). E, F. Hit3 and the positive control Ken rescued the ability of ALS-hiMNs to form neuromuscular junctions (NMJs, indicated by arrows) on co-cultured myotubes. NMJ frequencies were indicated as percentage values over 100 hiMN network-associated myotubes counted for each group. Scale bar, 10 µm.

We previously showed that ALS-hiMNs exhibited poor survival but also smaller soma and less complex neurites, all of which could be much improved by Ken (10). Such morphological improvements of ALS-hiMNs could also be observed for cells treated with Hit3 (Fig. 2C). Compared to the vehicle controls, Hit3 robustly increased neurite complexity via promoting outgrowth and branching of neuronal processes when examined at 52 days. Additionally, Hit3-treated ALS-hiMNs showed large somas, comparable to those treated with Ken (Fig. 2D).

To further examine the functional effect of Hit3, we employed a co-culture system including ALS-hiMNs and primary mouse skeletal myotubes on top of primary astrocytes (10). NMJs were detected by α-BTX staining, whereas hiMNs and skeletal muscles were revealed by staining of SYN1 and MHC, respectively (Fig. 2E, F). Under the vehicle control condition, both ALS1-hiMNs and ALS4-hiMNs rarely formed NMJs, consistent with our previous observation (10). Remarkably, Hit3-treated ALS-hiMNs could robustly form NMJs on the cocultured myotubes, comparable to those treated with the positive control Ken (Fig. 2E, F). Together, these data indicate that Hit3 could remarkably rescue both the morphological and functional deficits of ALS-hiMNs.

### MAP4Ks are major the targets of Hit3 for protecting ALS-hiMNs

Hit3 (K02288) was initially identified as a potent inhibitor of BMP/ALK signaling (31). To determine whether ALK inhibition was involved in ALS-hiMN protection, a list of 12 ALK inhibitors was further evaluated. While Hit3 consistently showed robust protection, the other ALK inhibitors including the widely used LDN-193189 and LDN-212854 had only minor or no protection (Fig. S1A), indicating that ALK inhibition might not be a mechanism for Hit3-mediated neuroprotection.

In addition to ALK inhibition, kinome-wide analysis showed that Hit3 also potently inhibits HGK (MAP4K4), MINK1 (MAP4K6) and TNIK (MAP4K7) (31). These three MAP4Ks belong to the GCK-IV family of the STE20 group kinases (32, 33). They share similar protein structures and exhibit very high homology within the kinase domain (>90% amino acid identity) (34). To examine a potential role of MAP4Ks in ALS-hiMNs, we first tested another chemical inhibitor, PF6260933 (designated as MAP4Ki), an HGK-specific inhibitor but also potently inhibiting MINK1 and TNIK (35). Interestingly, MAP4Ki treatments could also greatly improve survival of ALS1-hiMNs (Fig. S1B, C), resembling the effect of Hit3 and indicating that MAP4K inhibition might be an underlying mechanism for Hit3’s action.

We next took a genetic approach by knocking down each of the three MAP4Ks. sgRNA-mediated knockdowns were confirmed through western blotting analysis (Fig. 3A). These sgRNAs individually or in combinations were then tested on ALS1-hiMNs. While knocking down individual MAP4K had not much effect, their combinations especially triple knockdowns significantly improved survival of ALS1-hiMNs when compared to the control sgRNA (sgLacZ) at both 1 week and 3 weeks post co-culture on astrocytes (Fig. 3B, C). Neuronal survival was accompanied with enhanced neurite outgrowth and branching, as well as bigger somas after MAP4K triple knockdowns when compared to the control (Fig. S1D). Of note, sgRNAs were less effective than Hit3 treatment, which might be due to a lower efficiency of targeting these kinases through genetic knockdowns than chemical inhibition.

**Figure 3.**
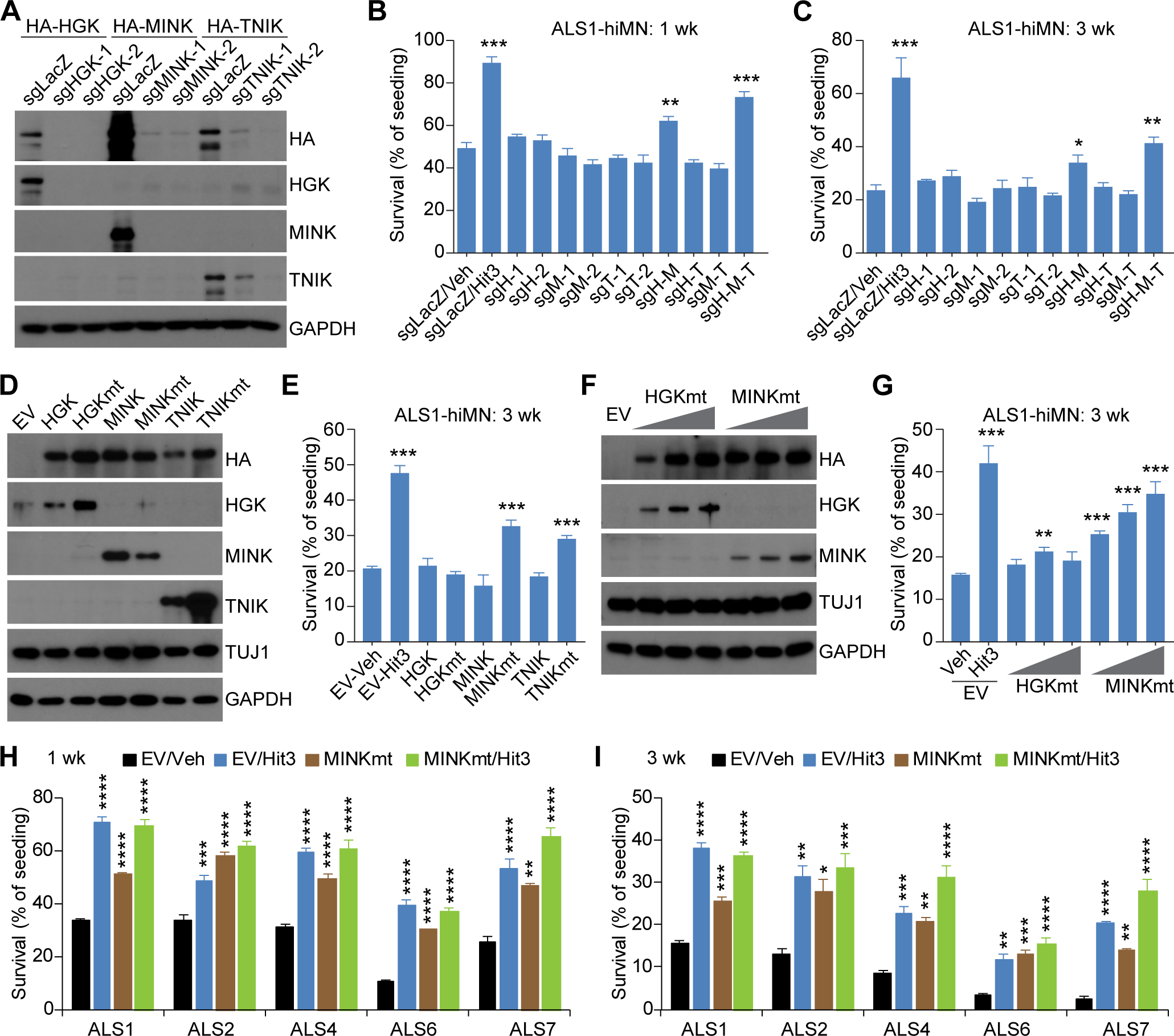
MAP4Ks are targets of Hit3 for improving ALS-hiMNs. A. Western-blotting analysis of knocking down MAP4Ks. HA-tagged MAP4Ks were co-expressed with sgRNAs (two for each kinase) and Cas9 in human fibroblasts and examined 10 days later. B, C. Knockdowns of MAP4Ks improve survival of ALS1-hiMNs, examined at 1 week or 3 weeks post replating on astrocytes (mean ± SEM; n = 4 independent samples; *p < 0.05, **p < 0.01, and ***p < 0.001 when compared to the control sgLacZ/Veh group, Student’s t-test). sgH, sgM, and sgT represent sgRNA against HGK, MINK1, and TNIK, respectively. D. Western-blotting analysis of ectopic MAP4Ks or their kinase-dead mutants in ALS1-hiMNs. TUJ1 and GAPDH are loading controls. E. Kinase-dead mutants of MAP4Ks improve survival of ALS1-hiMNs. EV, empty vector. (mean ± SEM; n = 4 independent samples; ***p < 0.001 when compared to the control EV-Veh group, Student’s t-test). F. Western-blotting analysis of ectopic MAP4K mutants in ALS1-hiMNs. EV, empty vector. G. MINK1 mutant dose-dependently improves survival of ALS1-hiMNs, comparable to Hit3 treatment (mean ± SEM; n = 4 independent samples; **p < 0.01 and ***p < 0.001 when compared to the control EV-Veh group, Student’s t-test). H, I. MINK1 mutant, comparable to Hit3, promotes survival of ALS-hiMNs from diverse human patients (mean ± SEM; n = 4 independent samples; *p < 0.05, **p < 0.01, ***p < 0.001 and ****p < 0.0001 when compared to the control EV-Veh group, Student’s t-test). Cells were examined at 1 week or 3 weeks after replating on astrocytes.

To further examine the role of MAP4Ks, we took a dominant-negative approach through ectopic expression of kinase-dead mutants, including HGK-K54R (HGKmt), MINK1-K54R (MINK1mt) and TNIK-K54R (TNIKmt) (36). These mutants and their wild type controls were introduced during reprogramming of patient fibroblasts to ALS1-hiMNs (Fig. 3D, E). Compared to the empty virus control, wild type kinases showed toxicity during the early reprogramming process, although they had not much effect once the reprogrammed ALS-hiMNs were replated onto astrocytes (Fig. 3E). Interestingly, the kinase-dead MINK1mt or TNIKmt improved the survival of ALS-hiMNs (Fig. 3E). Further analysis showed that MINK1mt dosage-dependently enhanced the survival of ALS-hiMNs, with an effect at the highest dosage comparable to those treated with Hit3 (Fig. 3F, G). In contrast, the HGKmt was less effective at all the examined dosages.

In comparison to Hit3, we examined MINK1mt on additional ALS-hiMNs including those derived from patients with mutations on FUS, SOD1, TDP43, and C9ORF72. MINK1mt and Hit3 showed comparable effect on survival of all these ALS-hiMNs, although the latter was slightly more effective potentially due to more efficient inhibition of MAP4Ks (Fig. 3H, I). Altogether, these results indicate that Hit3-mediated inhibition of MAP4Ks improves survival of ALS-hiMNs.

### CNH domain of MINK1 mimics the effect of Hit3

MINK1 consists of three highly conserved domains: a kinase domain, an intermediate domain, and a citron homology domain (CNH) (34). To map out the functional domain for MINK1mt-mediated neuroprotection, we constructed a series of HA-tagged truncations of MINK1mt (Fig. S2A). Expression of these constructs in ALS-hiMNs was confirmed by western blots (Fig. S2B). When examined at 1 week or 3 weeks post replating and compared to the empty virus controls, truncation mutants lacking the CNH domain had no effect (1-320, 1-959, and 296-959; Fig. S2C-F). In contrast, CNH-containing truncations (296-1312 and 866-1312) improved survival and morphology of ALS-hiMNs, comparable to those treated with Hit3 or full-length MINK1mt (Fig. S2C-G). Further truncation mutants showed that the flanking sequences surrounding the CNH domain were also important, as the mutant 994-1312 or 960-1292 largely failed to promote survival of ALS-hiMNs (Fig. S2E, F). Together, these results indicate that the CNH domain of MINK1 may play a dominant-negative function in improving ALS-hiMNs.

### MAP4Ks interact with RANGAP1 and affect its subcellular distribution

To determine how inhibition of MAP4Ks regulates ALS-hiMNs, we first examined the role of p38 and JNK since they were reported to be the downstream signaling targets and shown to be involved in neuronal degeneration (23, 24, 37, 38). However, their inhibition through a panel of small molecules appeared to be either non-effective or toxic to the survival of ALS1-hiMNs (Fig. S3A, B), suggesting that neither p38 nor JNK mediates the effect of MAP4Ks in ALS1-hiMNs.

We then took an unbiased approach to identify MAP4K-associated targets through BioID2-mediated protein proximity labeling (39). ALS1-hiMNs were transduced with lentivirus expressing either BioID2-MINK1mt or MINK1mt alone and treated with biotin. Biotinylated proteins were efficiently pulled down by streptavidin Dynabeads (Fig. 4A). Two batches of samples were analyzed by mass spectrometry. With a cutoff of 2.5-fold enrichment, 443 proteins were common to these two sets of data (Fig. 4B), including those known MINK1 interactors (Table S4), such as NCK2, STRN, STRN3, STRN4, HGK, TANC1, and TRIM25. These 443 proteins were subjected to KEGG pathway analysis. The top 5 pathways were Proteasome, Ribosome, Pathogenic E. coli Infection, RNA Transport, and Aminoacyl-tRNA Biosynthesis (Fig. 4C, D, and Table S4). Dysfunctions of proteasome, ribosome, RNA transport, and tRNA biogenesis are known to be associated with neurodegeneration including ALS (40, 41). Several tubulins are included in the Pathogenic E. coli Infection pathway, which involves in microtubule destabilization. Among these tubulins, TUBA4A mutations were recently identified in familial ALS (42). Similarly, several nuclear pore proteins in the RNA Transport pathway, including RANBP2, RANGAP1, and NUP214, are also closely involved in ALS (43–46). Such data further implicate MAP4Ks in the regulation of this type of neurodegeneration.

**Figure 4.**
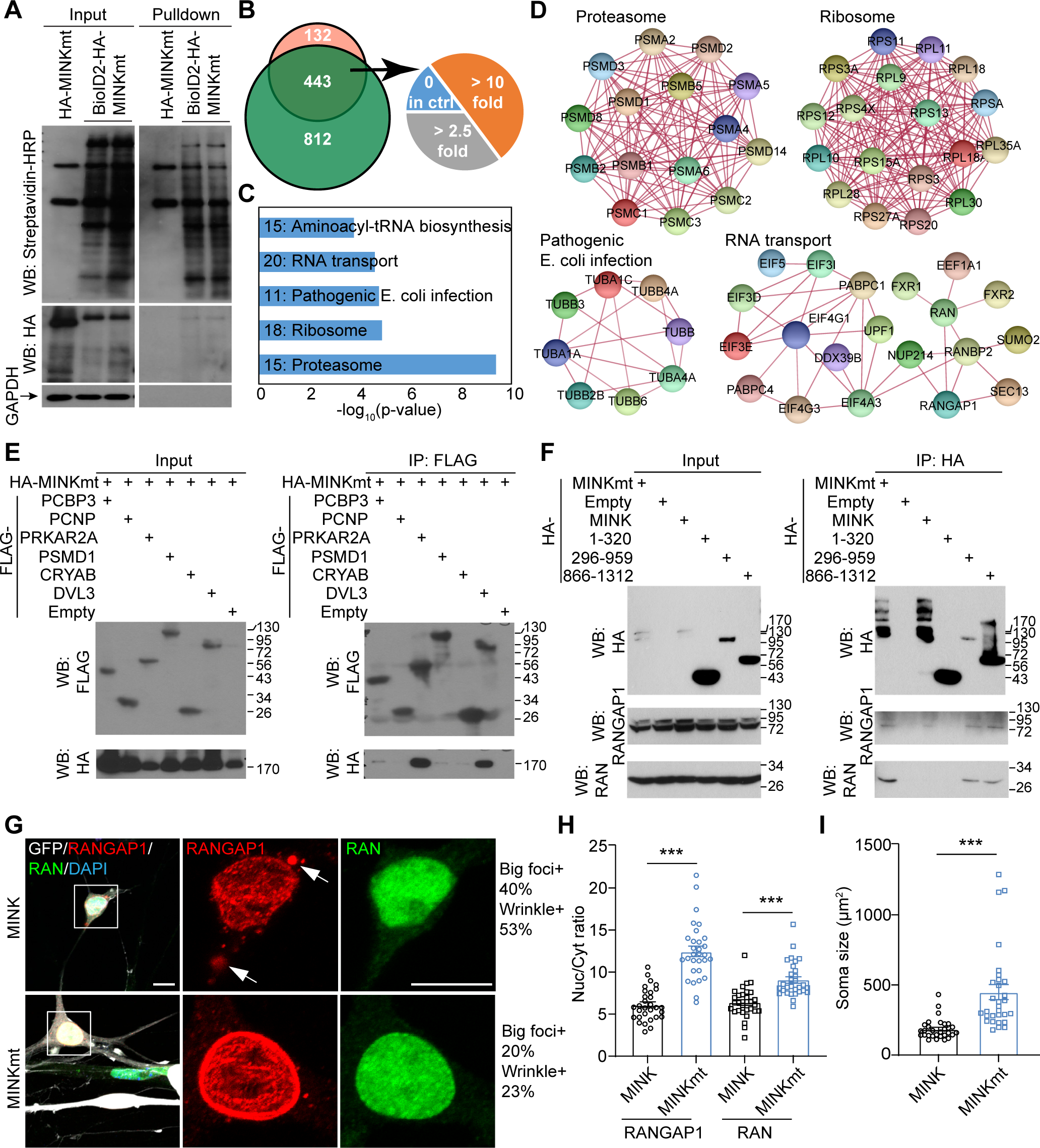
MAP4K interactome and the effect on RANGAP1 subcellular distribution. A. Western-blotting analysis of proteins after proximity labeling in ALS1-hiMNs at 10 dpi. B. Venn diagram of proteins with >2.5-fold enrichment in two biological repeats. C. The top 5 KEGG pathways. D. STRING analysis of association networks of proteins in the top 4 KEGG pathways. E. Validation of protein associations by co-immunoprecipitations (co-IP) and western blots. F. co-IP assays showing association of MINK1mt with RANGAP1 or RAN. G. Confocal images showing subcellular distribution of RANGAP1 or RAN in ALS1-hiMNs. Arrows indicate aggregated cytoplasmic RANGAP1 foci. Scale bar, 10 µm. H. MINKmt improves nuclear/cytoplasmic (Nuc/Cyt) ratios of the indicated proteins in ALS1-hiMNs (mean ± SEM; n = 30 neurons per group; ***p < 0.001, Student’s t-test). I. MINKmt improves soma sizes of ALS1-hiMNs (mean ± SEM; n = 30 neurons per group; ***p < 0.001, Student’s t-test).

We examined a few top candidate interactors through co-immunoprecipitation and western blotting. Both strong (PRKAR2A and DVL3) and weak (PCBP3, PSMD1, CRYAB) interactions could be identified (Fig. 4E), consistent with the principle of BioID2-mediated protein proximity labeling that identifies both interacting proteins and those in the vicinity (39). Co-immunoprecipitation assays also showed that MINK1 interacted with endogenous RANGAP1 and RAN (Fig. 4F). Domain mapping of MINK1 indicated that both its intermediate domain (aa296-959) and CNH domain (aa866-1312) could mediate interactions with RANGAP1 or RAN (Fig. 4F). Functionally, ectopic MINK1mt significantly increased the nuclear/cytoplasmic (Nuc/Cyt) ratio of RANGAP1 or RAN in ALS1-hiMNs (Fig. 4G, H). It also greatly promoted soma size of these neurons (Fig. 4I). Together, these results indicate that MAP4Ks regulate nuclear pore function and their inhibition protects ALS-hiMNs.

### MAP4K inhibition improves nuclear pore proteins and those involved in ALS

The above data from proteomics prompted us to examine the role of Hit3 on subcellular distribution of nucleopore proteins and those involved in ALS. We compared hiMNs from five ALS patients and three healthy controls. RANGAP1-containing cytoplasmic foci could be observed in all these hiMNs; however, quantifications showed that the Nuc/Cyt ratio of RANGAP1 was significantly lower in ALS-hiMNs than in the control hiMNs (Fig. 5A-C).

**Figure 5.**
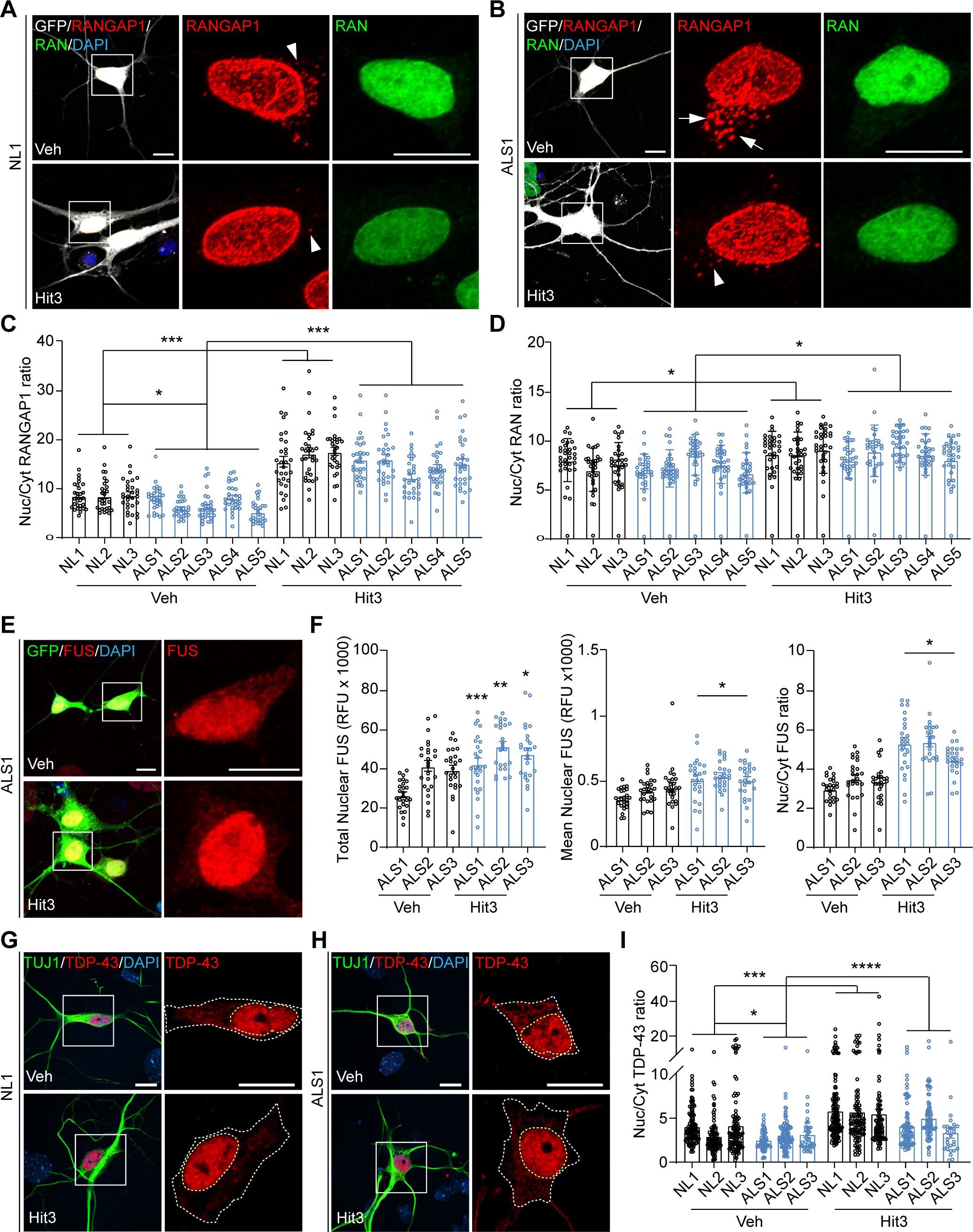
Hit3 improves nucleocytoplasmic transport in ALS-hiMNs. A, B. Confocal images of the indicated proteins in NL1-or ALS1-hiMNs. Cytoplasmic foci of RANGAP1 are indicated by arrows or arrowhead. Scale bar, 10 µm. C. Hit3 improves nuclear localization of RANGAP1 in hiMNs (n = 30 neurons per group; *p < 0.05 and ***p < 0.001, ordinary one-way ANOVA, comparisons between groups). D. Hit3 improves nuclear localization of RAN in hiMNs (n = 30 neurons per group; *p < 0.05, ordinary one-way ANOVA, comparisons between groups). E. Confocal images of FUS in ALS1-hiMNs. Scale bar, 10 µm. F. Hit3 improves nuclear fraction of FUS in ALS-hiMNs with different FUS mutations (n = 25 neurons per group; *p < 0.05, **p < 0.01 and ***p < 0.001, Student’s t-test for the left panel when compared to samples treated with Veh, one-way ANOVA for comparisons between groups in the middle and right panels). G, H. Confocal images of TDP-43 expression in NL1-or ALS1-hiMNs. Scale bar, 10 µm. I. Hit3 improves nuclear fraction of TDP-43 in ALS-hiMNs (n > 28 neurons per group; *p < 0.05, ***p < 0.001, and ****p < 0.0001, ordinary one-way ANOVA, comparisons between groups).

Interestingly, Hit3 treatments greatly improved the Nuc/Cyt ratio of RANGAP1 in all these hiMNs, with largely disappeared cytoplasmic RANGAP1-containing foci (Fig. 5A-C). Hit3 also markedly improved the Nuc/Cyt ratio of RAN, despite a lack of difference between NL- and ALS-hiMNs under the untreated condition (Fig. 5A, B, D).

We previously showed that FUS protein is mislocalized in the cytoplasm of ALS-hiMNs with mutated FUS gene (10). Hit3 treatments significantly improved its nuclear localization, indicated by the increased total nuclear protein and the Nuc/Cyt ratio of FUS (Fig. 5E, F). A lower Nuc/Cyt ratio of TDP-43 was also observed in ALS-hiMNs when compared to NL-hiMNs (Fig. 5G-I). Hit3 treatments similarly enhanced nuclear localization of TDP-43 in all these samples (Fig. 5I).

To confirm a role of MAP4Ks in these above functions of Hit3, we took a shRNA-mediated knockdown approach. Knockdown efficiency of HGK, MINK1, or TNIK was individually validated through qRT-PCR (Fig. S4A). These shRNAs and a control shRNA were then examined in ALS1-hiMNs. A mixture of shRNAs against all three MAP4Ks was most efficient in improving the Nuc/Cyt ratio of RANGAP1, with a result comparable to those Hit3-treated ALS1-hiMNs (Fig. S4B). This mixture of shRNAs also significantly increased the Nuc/Cyt ratio of RAN or TDP-43 (Fig. S4C). All together, these results indicate that Hit3, through inhibition of MAP4Ks, can significantly enhance nuclear localization of proteins controlling nucleopore function and those involved in ALS.

### MAP4Ks regulate RANGAP1 distribution through a HDAC6-TUBA4A axis

How is subcellular distribution of RANGAP1 regulated by MAP4Ks? Previous studies show that those cytoplasmic RANGAP1+ foci are pore complexes associated with annulate lamellae (AL), membrane sheets of the endoplasmic reticulum (47, 48). MAP4K-mediated phosphorylation could cause redistribution of RANGAP1 between AL and nuclear membrane. This was examined by Phos-tag gels and western blots with antibodies broadly recognizing phospho-serine (pSer) or phospho-threonine (pThr). Unexpectedly, neither HGK nor MINK1 induced phosphorylation changes of RANGAP1 (Fig. S5A).

It was previously shown that microtubule-dependent transport of AL-localized pore complexes regulate biogenesis of nuclear pores (48, 49). Interestingly, our proximity-labeling proteomics identified many tubulins that are associated with MAP4Ks (Fig. 4D). Among these, TUBA4A is especially interesting due to its genetic association with familial ALS (42). Co-immunoprecipitation assays confirmed that MAP4Ks and TUBA4A are within the same protein complex (Fig. S5B). Through an shRNA approach, we found that downregulation of TUBA4A significantly reduced nuclear localization of RANGAP1 in ALS1-hiMNs (Fig. 6A-C). Once again, however, we failed to detect MAP4K-induced phosphorylation of TUBA4A, indicating alternative regulation of TUBA4A by MAP4Ks (Fig. S5C-D).

**Figure 6.**
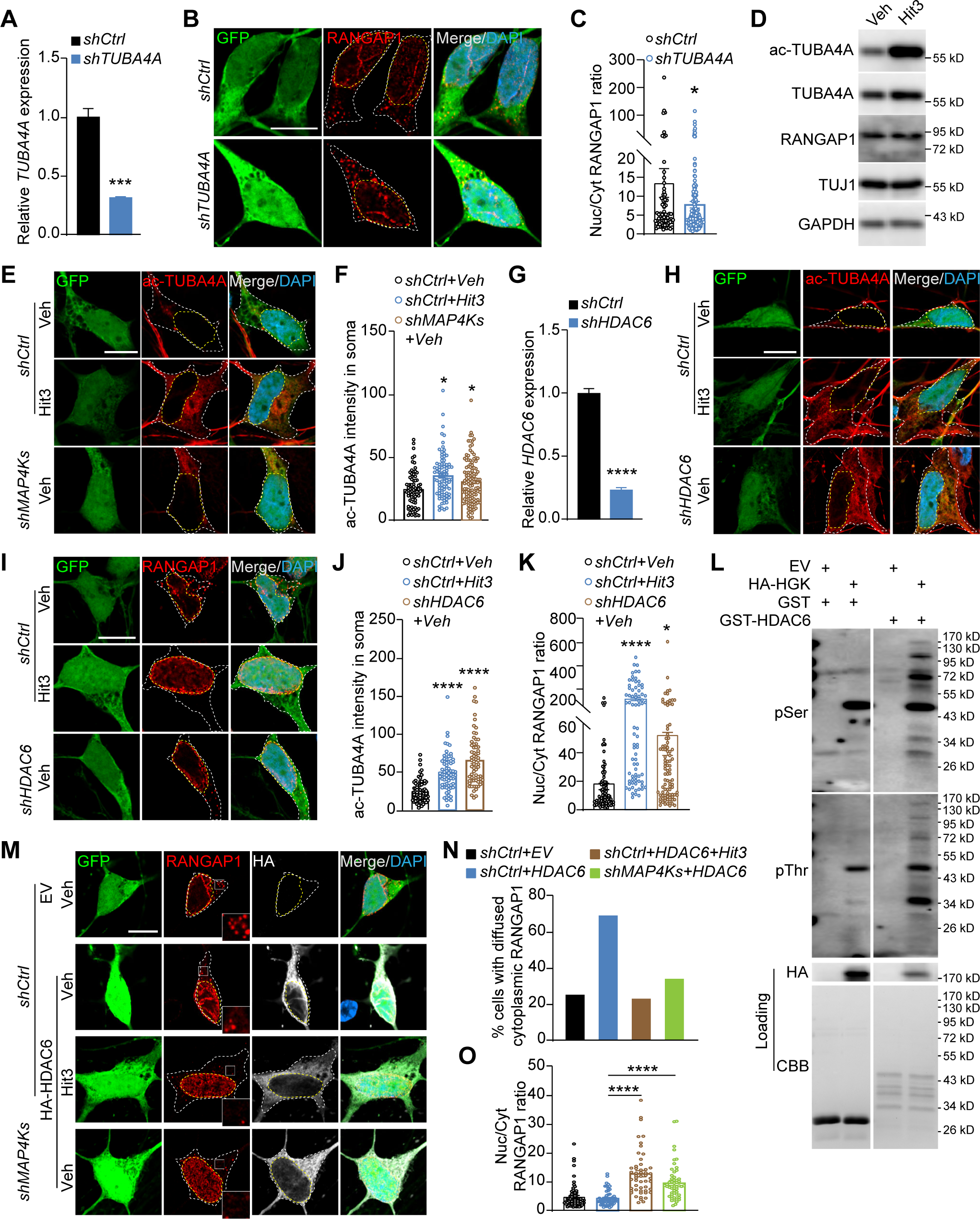
A role of the MAP4K-HDAC6-TUBA4A axis in subcellular distribution of RANGAP1. A. qRT-PCR analysis of shRNA-mediated knockdown of endogenous TUBA4A in human fibroblasts (mean ± SEM; ***p < 0.001, Student’s t-test). B. Confocal images of RANGAP1 distribution in ALS1-hiMNs co-cultured with astrocytes at 28 dpi. Scale bar, 10 µm. C. TUBA4A knockdown increases cytoplasmic fraction of RANGAP1 in ALS1-hiMNs (mean ± SEM; *p < 0.05, Student’s t-test). D. Western blots showing enhanced acetylation of TUBA4A (ac-TUBA4A) by Hit3 in ALS1-hiMNs. E. Confocal images of ac-TUBA4A in ALS1-hiMNs cocultured with astrocytes at 28 dpi. Scale bar, 10 µm. F. Knockdown of MAP4Ks, comparable to Hit3 treatments, enhances TUBA4A acetylation in somas of ALS1-hiMNs (mean ± SEM; *p < 0.05, ordinary one-way ANOVA, when compared to *shCtrl+Veh* group). G. qRT-PCR analysis of shRNA-mediated knockdown of endogenous HDAC6 in human fibroblasts (mean ± SEM; ****p < 0.0001, Student’s t-test). H. Confocal images of ac-TUBA4A in ALS1-hiMNs cocultured with astrocytes at 28 dpi. Scale bar, 50 µm. I. Confocal images of RANGAP1 distribution in ALS1-hiMNs cocultured with astrocytes at 28 dpi. Scale bar, 10 µm. J. Knockdown of HDAC6 promotes TUBA4A acetylation in somas of ALS1-hiMNs (mean ± SEM; ****p < 0.0001, ordinary one-way ANOVA, when compared to *shCtrl+Veh* group). K. Knockdown of HDAC6 promotes nuclear localization of RANGAP1 in ALS1-hiMNs (mean ± SEM; *p < 0.05 and ****p < 0.0001, ordinary one-way ANOVA, when compared to *shCtrl+Veh* group). L. In vitro kinase assay and western blotting showing phosphorylation of purified GST-HDAC6 by HGK. M. Confocal images of RANGAP1 distribution in ALS1-hiMNs cocultured with astrocytes at 28 dpi. Scale bar, 10 µm. N. Knockdown of MAP4Ks, as well as Hit3 treatments, reduces hiMNs with HDAC6-induced abnormal cytoplasmic distribution of RANGAP1. O. Knockdown of MAP4Ks, as well as Hit3 treatments, promotes nuclear localization of RANGAP1 even in the presence of HDAC6 (mean ± SEM; ****p < 0.0001, ordinary one-way ANOVA, when compared to *shCtrl+Veh* group).

Tubulin acetylation enhances microtubule stability and intracellular transport (50). We examined whether acetylation status of TUBA4A could be regulated by MAP4Ks. When cells were treated with Hit3, western blotting showed that acetylated TUBA4A (ac-TUBA4A) was greatly increased indicating a negative regulation by MAP4Ks (Fig. 6D). Immunocytochemistry also showed that the level of ac-TUBA4A was enhanced in the somas of ALS1-hiMNs when treated with Hit3 or after shRNA-mediated knockdown of endogenous MAP4Ks (Fig. 6E, F).

Tubulin acetylation is governed by two opposing enzymes: alpha-tubulin acetyltransferase 1 and histone deacetylase 6 (HDAC6). HDAC6 was previously shown to play a role in protein aggregation, axonal transport, and pathological phenotypes of ALS models (51–58). We then focused on HDAC6 and found that knockdown of HDAC6 significantly enhanced the levels of ac-TUBA4A in the somas of ALS1-hiMNs (Fig. 6G, H, J, S6A). Accordingly, HDAC6 knockdown markedly increased RANGAP1 localization to the nucleus (Fig. 6I, K), indicating an inverse correlation of HDAC6 activity and RANGAP1 nuclear localization. HDAC6 knockdown also promoted nuclear localization of TDP-43 (Fig. S6B, C), consistent with a previous report (56).

A motif scan revealed that HDAC6 harbors five predicted MAP4K phosphorylation sites (59). We then performed a phosphorylation assay by using purified GST-HDAC6 as the substrate and immunoprecipitated HA-HGK as the kinase. Both full-length and some degraded forms of GST-HDAC6 could be phosphorylated by HGK when examined by western blotting with antibodies for pSer or pThr (Fig. 6L). Together with the findings showing an increased level of ac-TUBA4A when MAP4Ks or HDAC6 is inhibited (Fig. 6F, J), these results indicate that MAP4K-mediated phosphorylation may enhance HDAC6 activity. Such a conclusion will be consistent with previous reports demonstrating that the alpha-tubulin deacetylase activity of HDAC6 is stimulated through phosphorylation by multiple kinases (60–62).

To further determine the functional relationship of MAP4Ks and HDAC6, we overexpressed HDAC6 in ALS1-hiMNs with or without Hit3 treatments or MAP4K knockdowns. Ectopic HDAC6 resulted in a higher percentage of cells with diffused cytoplasmic RANGAP1 and a much-reduced Nuc/Cyt ratio of TDP-43 (Fig. 6M-O, S6D, E). Such cellular phenotypes could be reversed by Hit3 treatments or MAP4K knockdowns, indicating requirements of endogenous MAP4Ks (Fig. 6M-O, S6D, E). Collectively, our results suggest that MAP4Ks phosphorylate and stimulate HDAC6 activity, which leads to a reduction of ac-TUBA4A and the nuclear pore-localized RANGAP1.

### Inhibition of MAP4Ks is neuroprotective in vivo

To examine a potential role of MAP4K inhibition in vivo, we used the SOD1^G93A^ mouse model of ALS (63, 64). Pharmacokinetics showed that the MAP4K inhibitor PF6260933 (MAP4Ki) exhibited higher concentrations than Hit3 in both the plasma and the central nervous system after intraperitoneal injections (Table S5). As such, we used the MAP4Ki for subsequent in vivo studies. To further increase drug accessibility to the nervous system, we performed intrathecal injections starting at 40 days of age and until the end stage. When compared to the vehicle control, MAP4Ki had no significant effects on either body weights or performances on rotarod tests (Fig. 7A, B). Very interestingly, Kaplan-Meier survival curves showed that MAP4Ki significantly prolonged the median lifespan of SOD1^G93A^ mice (Fig. 7C; median survival: vehicle = 129 d, MAP4Ki = 139 d; χ^2^ = 8.975, **p = 0.0027, log-rank Mantel-Cox test). Such a beneficial effect of MAP4K inhibition closely mimics that of *Hdac6* deletion in the SOD1^G93A^ mice (52).

**Figure 7.**
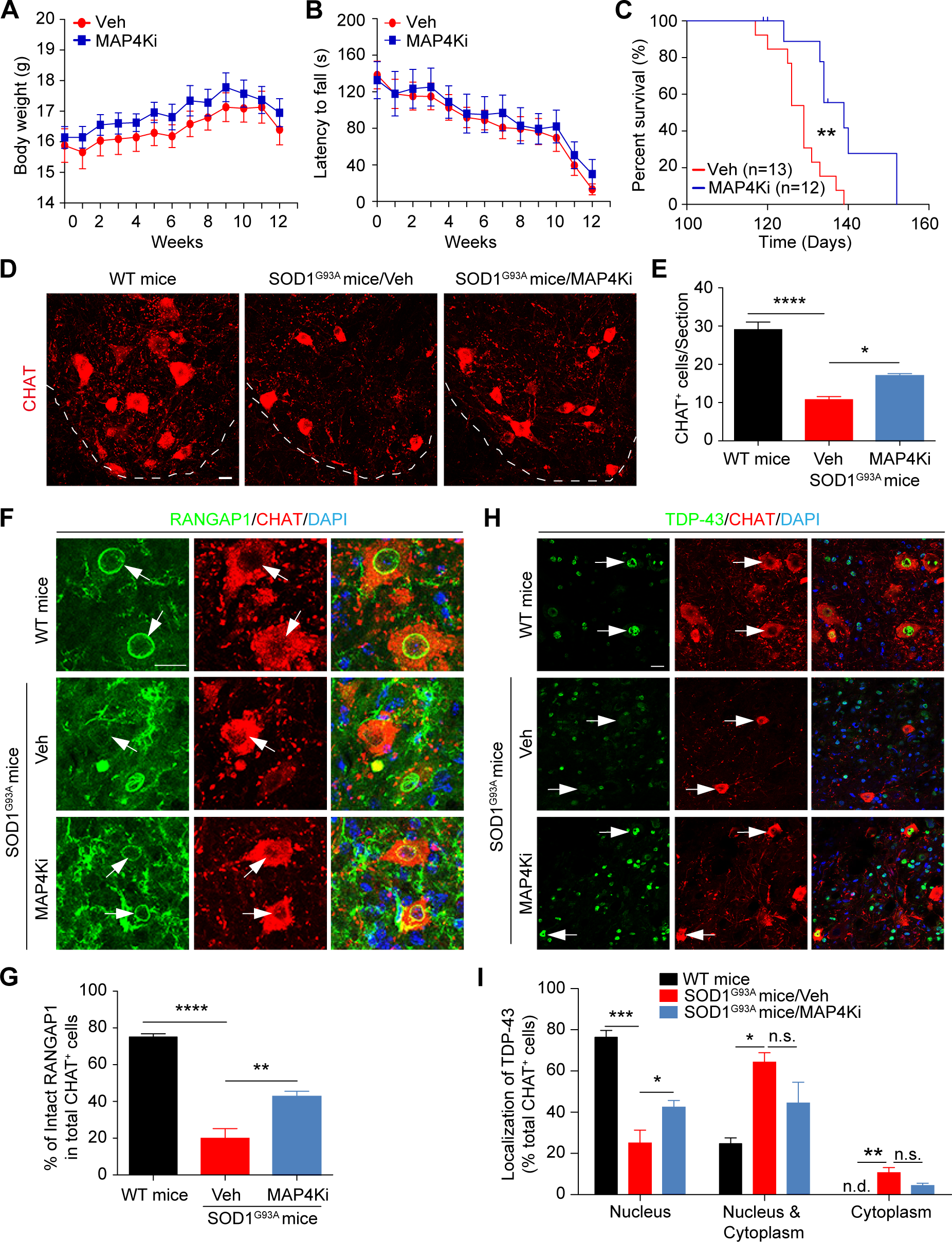
MAP4Ki is neuroprotective in SOD1^G93A^ mice. A. Body weight changes (n = 12-13 female mice per group). B. Rotarod tests (n = 12-13 female mice per group). C. Kaplan-Meier survival curve. Mice treated with MAP4Ki survived significantly longer than mice with the vehicle control (mean ± SEM; n = 12-13 female mice per group; Median survival: Veh = 129 days and MAP4Ki = 139 days; **p = 0.0027 by log-rank Mantel-Cox test). D. Confocal images of CHAT+ neurons in the ventral horn of the lumbar spinal cord. Scale bar, 20 μm. E. MAP4Ki preserves more CHAT+ neurons (mean ± SEM; n = 4-5 mice per group; *p < 0.05 and ***p < 0.001, one-way ANOVA). F. Confocal images of RANGAP1 localization in CHAT+ neurons. Scale bar, 20μm. G. MAP4Ki preserves CHAT+ neurons with nuclear membrane localized RANGAP1 (mean ± SEM; n = 3-4 mice per group; **p < 0.01 and ****p < 0.0001, one-way ANOVA). H. Confocal images of TDP-43 localization in CHAT+ neurons. Scale bar, 20μm. I. MAP4Ki preserves CHAT+ neurons with nuclear TDP-43 (mean ± SEM; n = 3-5 mice per group; *p < 0.05, **p < 0.01, and ***p < 0.001, one-way ANOVA; n.s., not significant; n.d., not detected).

The effect of MAP4Ki on motor neurons was examined through immunohistochemistry by using another cohort of mice at about 126 days of age, a time point around the median survival of vehicle-treated mice. A dramatic reduction of CHAT+ motor neurons was observed in the ventral horns of the lumbar spinal cord of SOD1^G93A^ mice when compared to their wildtype counterparts (Fig. 7D, E; group effect: F(2, 11) = 47.61, ****p < 0.0001, one-way ANOVA), consistent with known pathological features of this ALS mouse model. Importantly, more CHAT+ motor neurons were preserved in mice treated with MAP4Ki than the vehicle control (Fig. 7D, E; *p = 0.0206, one-way ANOVA and Tukey’s post hoc test).

RANGAP1 expression in motor neurons was examined by immunostaining. It was mainly localized in the ring-like nuclear membrane in CHAT+ cells of wildtype mice (Fig. 7F). Such a localization was rarely observed in survived CHAT+ motor neurons of vehicle-treated SOD1^G93A^ mice. Interestingly, MAP4Ki treatments maintained a significantly larger number of motor neurons with clear RANGAP1 nuclear membrane expression (Fig. 7F, G; group effect: F(2, 7) = 57.46, ****p < 0.0001; MAP4Ki vs. Veh: **p = 0.0055; one-way ANOVA and Tukey’s post hoc test). We also examined TDP-43 expression in CHAT+ cells in the ventral horn of the lumbar spinal cord. It was predominantly detected in the nucleus in wildtype mice, whereas a large fraction of CHAT+ cells showed cytoplasmic localization of TDP-43 in vehicle-treated SOD1^G93A^ mice (Fig. 7H, I; group effect: F(2, 9) = 27.82, ***p = 0.0001, one-way ANOVA). In contrast, a significant number of CHAT+ cells maintained nuclear TDP-43 in SOD1^G93A^ mice when treated with MAP4Ki (Fig. 7H, I; *p = 0.0452, one-way ANOVA and Tukey’s post hoc test). On the other hand, MAP4Ki treatments had no effect on reactive gliosis, indicated by the expression of GFAP or IBA1 (Fig. S7). Together, these results show that inhibition of MAP4Ks is neuroprotective and prolongs survival of SOD1^G93A^ mice, potentially mediated through improved subcellular localization of proteins involved in motor neuron function.

## DISCUSSION

Our proof-of-concept high-throughput screens identify a lead compound, Hit3, which can significantly promote survival, growth, and function of aging-relevant ALS-hiMNs. Mechanistically, our results demonstrate that Hit3 mainly functions as an inhibitor of the GCK-IV subfamily of MAP4Ks. We further show that the MAP4Ks-HDAC6-TUBA4A pathway regulates the subcellular distribution of RANGAP1, a key component of the nuclear pore complexes (NPCs), and TDP-43, an RNA/DNA binding protein involved in human ALS. Importantly, MAP4Ki treatment of SOD1^G93A^ mice preserves motor neurons and extends animal survival.

This is the first to apply aging-relevant human neurons in screens for therapeutic small molecules. Unlike iPSC-derived neurons with age-rejuvenation (15, 18), directly reprogrammed neurons from skin fibroblasts of adult human patients reserve aging-associate features and are therefore advantageous in modeling adult-onset neurodegenerative diseases (13–15, 65–67). However, these aging-relevant human neurons have not been previously utilized for high-throughput chemical screens, partly due to limited neuronal purity and yield. With improved conversion efficiency and simple enrichment procedures, our current study shows the feasibility of employing these neurons in screens for small molecules and pathways that may be therapeutically relevant. Such a screening strategy may be applied to studying other neurodegenerative diseases such as Parkinson’s and Alzheimer’s disease by using directly reprogrammed human neurons.

Our results showed that MAP4Ks are the major targets of the lead compound Hit3. Hit3, formally known as K02288, was initially discovered as a selective inhibitor of the bone morphogenetic protein (BMP) type I receptor kinases (31). Nonetheless, our examination of multiple additional inhibitors targeting the BMP/TGFβ pathway failed to show a comparable effect on promoting survival of ALS-hiMNs, suggesting alternative pathways targeted by Hit3 in these neurons. Indeed, kinome-wide analysis clearly showed that Hit3 also potently inhibits the GCK-IV family of the STE20 group kinases (33), including HGK (MAP4K4), MINK1 (MAP4K6) and TNIK (MAP4K7) (31). Accordingly, PF6260933, a structural analog of Hit3 and a selective inhibitor of HGK as well as MINK1 and TNIK (35), exhibits a similar protective effect on ALS-hiMNs. Our individual and combinatorial knockdown experiments also confirm a critical and redundant role of these three highly homologous kinases in neuronal survival. Interestingly, the kinase-dead MINK1 or its CNH domain can function as a dominant-negative mutant improving survival of hiMNs that are derived from diverse ALS patients. Our conclusion on the role of MAP4Ks are consistent with previous studies implicating MAP4Ks in neural degeneration (23, 24, 38). One major difference is that HGK alone was identified to regulate death of iPSC-derived motor neurons under stress (23), a result that may reflect the differential response of embryonic versus aging-relevant neurons.

A key downstream effect of MAP4Ks, we revealed, is the subcellular localization of RANGAP1. Previously, the JNK signaling pathway was identified as a target of MAP4Ks in controlling degeneration and apoptosis of embryonic or iPSC-derived neurons under stress (23, 24, 38). However, its inhibition with multiple specific inhibitors is either toxic or non-effective to ALS-hiMNs, once again revealing a differential response of embryonic versus aging-relevant neurons. In contrast, our unbiased proteomics uncovers that MAP4Ks may regulate multiple biological processes, such as proteosome, ribosome, and RNA transport. Dysfunction of these processes is well-known to be involved in neurodegeneration including ALS (40, 41, 43–46, 68). We focused on RANGAP1 due to its abnormal subcellular distribution in ALS-hiMNs and that such abnormality can be significantly improved by either Hit3 or a dominant-negative MINK1 mutant. RANGAP1, a core component of the NPCs, is the GTPase converting RAN-GTP to RAN-GDP, an essential step for RAN-mediated nuclear import and export of proteins and RNAs (69). Mislocalization of RANGAP1 and RAN were previously reported in mouse primary MNs, cortical neurons and lymphoblast cells when the ALS-associated mutant proteins were expressed (46, 70), as well as in human Huntington iPSC-derived striatal neurons neurons (71–73) and ventral nerve cord cells after traumatic injury (74). Such mislocalizations may finally lead to dysfunctional nucleocytoplasmic transport (46, 70–75). Although our co-immunoprecipitation assays indicate that MAP4Ks and RANGAP1 are in the same protein complex, multiple efforts failed to show RANGAP1 being phosphorylated by MAP4Ks, indicating alternative regulation of RANGAP1 subcellular localization.

RANGAP1-associated pore proteins not only reside within the nuclear membrane but are also found in annulate lamellae (AL), membrane sheets of the rough endoplasmic reticulum (76). AL are highly abundant in neurons (48, 77) and are the storage compartment for nuclear pore proteins that can later be transported to the NPCs (48). Such transport requires stable microtubules since their disruption leads to increased levels of AL and AL-localized nuclear pore proteins (48, 78, 79). This is consistent with our data showing enhanced RANGAP1 localization in cytoplasmic foci after down-regulation of the MAP4K-associated tubulin TUBA4A, mutations of which can destabilize the microtubule network and are implicated in familial ALS (42). Instead of a direct regulation of TUBA4A through phosphorylation, our results rather indicate that MAP4Ks inhibit TUBA4A acetylation, a posttranslational modification that increases microtubule stability (80, 81). This is likely due to altered HDAC6 activity through phosphorylation by MAP4Ks, as indicated by our in vitro kinase assays.

HDAC6 is the major α-tubulin deacetylase in the nervous system (82). Its activity is regulated by multiple posttranslational modifications including phosphorylation. Phosphorylation of HDAC6 can increase its enzymatic activity or protein stability (60–62, 83, 84). MAP4Ks may also regulate HDAC6 in a similar fashion since their inhibition by Hit3 or shRNA-mediated knockdowns can promote acetylation of TUBA4A. At the functional level, our results demonstrate that HDAC6 downregulation enhances nuclear localization of RANGAP1 and TDP-43 in ALS-hiMNs. These results are consistent with previous studies showing that HDAC6 inhibition restores subcellular mislocalization of TDP-43 and improves NMJ morphology in iPSC-derived MNs (56, 58). Further supporting a functional interaction of MAP4Ks and HDAC6 is that either MAP4Ki (this study) or *Hdac6* deletion (52) can similarly promote survival of the SOD1^G93A^ ALS mice.

In summary, by using aging-relevant human neurons our chemical screens identified a neuroprotective chemical compound and MAP4Ks as potential therapeutic targets for treating ALS and potentially other neurodegenerative diseases. Our studies also reveal redundant functions of these highly homologous kinases and a novel molecular pathway by which they control neuronal degeneration. Future studies are warranted to develop more potent blood-brain barrier-penetrating MAP4K inhibitors as therapeutics for ALS and other neurodegenerative diseases.

## MATERIALS AND METHODS

### Animals

C57BL/6J (Jax #000664), B6SJLF1 (Jax #100012), and SOD1^G93A^ (Jax #002726) were purchased from the Jackson Laboratory. The SOD1^G93A^ strain was maintained by breeding male hemizygous carriers to B6SJLF1 hybrids. Standard PCRs as described by the Jackson laboratory were used for genotyping. All mice were housed under a controlled temperature and a 12-h light/dark cycle with free access to water and food in a barrier animal facility. Sample sizes were empirically determined. Animal procedures and protocols were approved by the Institutional Animal Care and Use Committee at UT Southwestern.

### Human fibroblasts

The human fibroblast line C9-4 and C9-5 were gifts of Dr. Corey lab (85). Other fibroblast lines were obtained from Coriell, ATCC, or Cedars-Sinai. All fibroblast lines with details were listed in Table S1. They were maintained in DMEM-high glucose supplemented with 15% fetal bovine serum and 1% penicillin/streptomycin at 37LJ and 5% CO_2_.

### Chemicals

The L1700 bioactive compound library was purchased from Selleck Chemicals. Kenpaullone (Ken) and A83-1 were obtained from Tocris. Forskolin (FSK), LDN193189.2HCl (LDN), PF6260933 (MAP4Ki), and other individual chemicals were ordered from Sigma or Selleck.

### Plasmids and virus production

Lentiviral plasmids for hiMNs were previously reported (10) and are available from Addgene (#90214 and #90215). cDNAs for HA-tagged HGK, MINK1, TNIK, and their kinase-dead mutants were individually subcloned into a third-generation lentiviral vector, pCSC-SP-PW-IRES-GFP. This vector was also used to express BioID2-myc-HA-MINK1mt. A single vector CRISPR/Cas9 system was developed by replacing CMV-SP-PW-IRES-GFP with the hU6-Filler-EFS-SpCas9-FLAG-P2A-Puro cassette from lentiCRISPRv2 (Addgene #52961). To monitor the transduction rate, mCherry was inserted to replace the Puro fragment to construct pCSC-hU6-Filler-EFS-SpCas9-FLAG-P2A-mCherry. Then each individual sgRNA was subcloned into this new vector by replacing the Filler fragment. All primers used for PCR and/or subcloning of cDNA or sgRNA are listed in Table S6. Replication-incompetent lentiviruses were generated in HEK293T cells (ATCC) via co-transfections of lentiviral vectors, pREV, pMDL and pVSV-G. They were stored at 4LJ before cell transductions.

### Fibroblast-derived human induced motor neurons (hiMNs)

NL- and ALS-hiMNs were converted from adult human skin fibroblasts as previously described with modifications (10). Briefly, fibroblasts were plated onto Matrigel-coated 10-cm dishes at 1.5×10^4^ cells per cm^2^. The next day cells were transduced with lentiviral supernatants containing 6 µg/ml polybrene. After overnight transduction, fibroblasts were cultured in fresh fibroblast culture media for one more day. They were then switched into C2 medium consisting of DMEM:F12:neurobasal (2:2:1), 0.8% N2 (Invitrogen), 0.8% B27 (Invitrogen), and supplemented with 10 µM FSK, 0.5 µM LDN, and 10 ng/ml FGF2 (PeproTech). Medium was half-changed every other day until replating at 14 days post virus infection (dpi). The replating procedure was performed to isolate hiMNs from nonconverted fibroblasts (10). hiMNs were further enriched by passing through a 20-µm cell strainer and were then seeded into culture vessels coated with Matrigel or co-cultured with primary mouse cortical astrocytes, using C2 medium supplemented with 5 µM FSK and 10 ng/ml each of BDNF, GDNF, and NT3 (PeproTech). The medium was half-changed weekly until further analysis.

### Chemical screens

For primary and secondary screens, hiMNs were plated into Matrigel-coated 96-well plates. They quickly attached to the surface and outgrew processes within 2-4 hours. Individual chemical was then added into each well at 2.5 µM for primary screens and at 0.5, 1.0, 2.5, or 5 µM for secondary screens. Vehicle (DMSO) and Ken in quadruplicates served as the negative and positive control, respectively. Viable cells were then determined by the CellTiter-Glo Luminescent Cell Viability Assay. As previously reported (86), assay quality for each plate was evaluated by the Z-prime value [=1-3*((STDEV_Ken_+STDEV_Veh_)/ABS(Mean_Ken_-Mean_Veh_))], whereas relative survival for each chemical was calculated by (Sample-Mean_Veh_)/((Mean_Ken_-Mean_Veh_)*100). Chemicals were then ranked based on their effects on relative survival of ALS-hiMNs. Top hits with a greater than 1.5x standard deviations above the mean in primary screens were selected for secondary screens and those with greater than 3x standard deviations above the mean in secondary screens were selected for subsequent studies. All top hits from secondary screens were further assessed in dose-response assays (7-points in triplicates). Relative survival for each dosage was calculated via normalization to the vehicle controls.

For long-term chemical treatments, hiMNs were co-cultured with mouse primary cortical astrocytes seeded onto 96-well plates or coverslips in 24-well plates. One 96-well plate was used to determine the number of seeded GFP+ hiMNs 4-hour post plating. The rest plates were treated with the top 15 hits, vehicle, Ken, and other selected chemicals in pentaplicates at a proper concentration determined by the dose-response assays. Chemical treatments were repeated weekly until analysis at 1, 3, or 5 weeks later. hiMNs were then fixed and stained with antibodies for GFP and the neuronal marker TUJ1. Cells in each well were imaged and quantified with a Cytation3 imaging reader and software (BioTek). As all GFP+ cells were also TUJ1+, viable neurons in each well were further confirmed by manual counting GFP+ cells. Relative survival was calculated by first normalizing to the number of seeded GFP+ cells, followed by normalization to the vehicle control.

### Primary astrocytes

Primary astrocytes were prepared from the cerebral cortices of postnatal day (P)1-P3 mouse pups as previously described(10). Contaminating neurons and microglia were removed via vigorous shaking and a few cycles of passaging, freezing, thawing, and replating. For co-culture with neurons, proliferating astrocytes were inhibited via treatments with 2 µM Ara-C, a mitotic inhibitor, for at least 48 hours.

### Neuromuscular junctions

Primary myoblasts were isolated from skeletal muscles of P0.5 mouse pups and differentiated into myotubes as previously described (10). Myotubes were resuspended in neuronal culture medium and then plated onto coverslips with co-cultured astrocytes and hiMNs at 40 to 50 dpi. After another 4 to 7 days, these sandwich cultures of myotubes, hiMNs, and astrocytes were live-stained with rhodamine-conjugated α-BTX (Invitrogen, 1:10,000) for 1 hour at 37LJ. α-BTX labelled cells were then processed for immunostaining with antibodies of SYN1 (Cell Signaling Technology, 1:500) and MHC (Sigma, 1:1,000). NMJ formation frequency was presented as the percentage of NMJs on myotubes associated with hiMNs networks.

### Immunocytochemistry

Cells were processed for immunocytochemistry as previously described (10). Antibodies used in this work were listed in Table S7. Nuclei were counterstained with Hoechst 33342 or DAPI. Images were obtained with a NIKON A1R confocal microscope, a Cytation3 imaging reader, or an EVOS fluorescence microscope. Confocal images were used for quantification of subcellular distribution of RANGAP1, RAN, TDP-43, or FUS, as well as the number of RANGAP1+ foci and NMJ+ myotubes. Fluorescence intensity was quantified as previously described with minor modifications (10). In brief, ImageJ with a plugin of Bio-Formats was used to measure total fluorescence intensity separately in the soma and the nucleus. Cytoplasmic intensity was obtained by subtracting the intensity in the nucleus from that in the soma. Subcellular distribution was then represented by the nuclear/cytoplasmic ratio.

### Gene overexpression or knockdown in hiMNs

Lentivirus carrying cDNA or sgRNA/Cas9 was co-transduced with the reprogramming lentiviruses in fibroblasts, using empty vector (EV) or sgLacZ/Cas9 as the respective control. After neuronal induction, the replated hiMNs at 14 dpi were directly used for western blotting or seeded onto mouse astrocytes-coated 96-well plates or coverslips in 24-well plates. The seeding density of GFP+ hiMNs in 96-well plates was determined 4-hour post plating as described above. These cultures were then processed for immunocytochemistry and analyzed for survival, morphology, soma size, and other features at the indicated time-points.

Doxycycline (Dox) inducible system was used for shRNA-mediated knockdowns. To determine knockdown efficiency, the lentiviruses FUW-M2rtTA and TRE3G-miRE-shRNA were applied to cultured fibroblasts 2∼6 hours after seeding. shRNA expression was induced by daily addition of Dox (0.5 µg/ml) into the medium. Cells were collected for qRT-PCR or western blotting 4 days post virus transduction. For knockdowns in hiMNs, the lentiviruses FUW-M2rtTA and TRE3G-miRE-shRNA were combined with the reprogramming factors. Cells were replated onto astrocyte-coated and Matrigel-treated coverslips. Dox (0.5 µg/ml) was added daily for the first week after replating and weekly thereafter. Hit3 (10 µM) was added twice in the first week after replating, then once per week. At the indicated time-points, cells were fixed for immunocytochemistry. Images of single confocal plane across the center of the nucleus were obtained on the NIKON A1R confocal microscope under a 100x objective with a pinhole setting at 1.2 under 488 nm laser. Because of the complexity of neuron-astrocyte co-cultures, neuronal nucleus and soma were manually defined by using the ImageJ program. The mean fluorescence intensity of acetylated TUBA4A was measured in the cytoplasm of hiMNs. The mean fluorescence intensity of RANGAP1, RAN, TDP-43 were separately measured in the cytoplasm or nucleus of hiMNs. The Nuc/Cyt ratios of different proteins were calculated by Microsoft Excel and analyzed by GraphPad Prism 9.

### Co-immunoprecipitation

Co-immunoprecipitation experiments were carried out as previously described with modifications (87). Briefly, cells were harvested and lysed in five volumes of lysis buffer (10 mM Tris-HCl, 150 mM NaCl, 1% NP-40, 1% Triton X-100, 1 mM dithiothreitol, 1% protein inhibitors (Pierce), 1x PhosSTOP (Roche), pH 7.5) for 20 min with rotations at 4LJ. Cell lysates were cleared by centrifugation at 21,000 g for 30 min. Equal amounts of proteins were incubated overnight with the anti-FLAG M2 (Sigma M8823) or anti-HA (Pierce #88836) magnetic beads at 4LJ. The beads were then sequentially washed for 3 min with gentle shaking with the following buffers: buffer 1 (10 mM Tris-HCl, 150 mM NaCl, 1% NP-40, pH 7.4), buffer 2 (10 mM Tris-HCl, 500 mM NaCl, pH 7.4), buffer 3 (10 mM Tris-HCl, 150 mM NaCl, pH 7.4), and buffer 4 (10 mM Tris-HCl, pH 7.4). Bead-bound proteins were eluted with 50 µl of 2 x SDS loading buffer, or twice (5 min each) of 20 µl of 0.1 M Glycine-HCl buffer (pH 3.0). The elutes were balanced by adding 10 µl solution consisting of 0.5 M Tris-HCl, 1.5 M NaCl, pH 7.4.

### Western blotting

Cells were collected in lysis buffer (50 mM Tris-HCl (pH 7.5), 150 mM NaCl, 1% Triton X-100, 0.1% SDS, 1% sodium deoxycholate, 0.01% NaN3, protease inhibitors (Pierce), and Phos-STOP (Roche). Protein concentration was determined by the Bradford assay. Protein samples were separated by SDS-PAGE and transferred to PVDF membranes. After blocking in 5% nonfat milk in PBST for 1 h at RT, membranes were incubated overnight with primary antibodies at 4 °C.

Horseradish peroxidase-conjugated secondary antibodies (Jackson) were applied, and the blots were developed with Pierce ECL Western Blotting Substrate (Thermo-Fisher) or Immobilon Western Chemiluminescent HRP substrate (Millipore-Sigma). Antibodies are listed in Table S7.

### Proximity-labeling proteomics

This was carried out according to a previously described protocol with modifications (88). Lentivirus expressing BioID2-myc-HA-MINK1mt or HA-MINK1mt was introduced during fibroblast reprogramming. After neuronal induction, the replated hiMNs at 10 dpi were seeded onto Matrigel-coated 6-well plates in neuronal culture medium containing 50 µM biotin. After 24 hours, biotin-treated cells were collected into lysis buffer composed of 50 mM Tris-HCl (pH 7.4), 500 mM NaCl, 0.2% SDS, and protease inhibitors (Pierce). Biotinylated proteins were pulled down with streptavidin beads following a published procedure (88). Samples were run 5∼10 mm into a 4 - 15% gradient precast protein gel and stained with G250 Coomassie brilliant blue. Each gel lane with protein bands was cut out and chopped into ∼1 mm^3^ cubes. Proteins were in-gel digested overnight with trypsin (Pierce), followed by reduction and alkylation with DTT and iodoacetamide (Sigma–Aldrich). Samples were then undergone solid-phase extraction cleanup with an Oasis HLB plate (Waters) and the resulting samples were injected onto an Orbitrap Fusion Lumos mass spectrometer coupled to an Ultimate 3000 RSLC-Nano liquid chromatography system. Samples were injected onto a 75 µm i.d., 75-cm long EasySpray column (Thermo) and eluted with a gradient from 0-28% buffer B over 90 min. Buffer A contained 2% (v/v) ACN and 0.1% formic acid in water, and buffer B contained 80% (v/v) ACN, 10% (v/v) trifluoroethanol, and 0.1% formic acid in water. The mass spectrometer operated in positive ion mode with a source voltage of 1.8 kV and an ion transfer tube temperature of 275 LJ. MS scans were acquired at 120,000 resolution in the Orbitrap and up to 10 MS/MS spectra were obtained in the ion trap for each full spectrum acquired using higher-energy collisional dissociation (HCD) for ions with charges 2-7. Dynamic exclusion was set for 25 s after an ion was selected for fragmentation. Raw MS data were analyzed using Proteome Discoverer v2.2 (Thermo), with peptide identification performed using Sequest HT searching against the human protein database from UniProt. Fragment and precursor tolerances of 10 ppm and 0.6 Da were specified, and three missed cleavages were allowed. Carbamidomethylation of Cys was set as a fixed modification, with oxidation of Met set as a variable modification. The false-discovery rate (FDR) cutoff was 1% for all peptides.

### Quantitative RT-PCR (qRT-PCR)

qRT-PCR was conducted as previously described with modifications (7). After removing medium, cells were collected in TRIzol reagent (Invitrogen) and total RNA was isolated by using a commercial kit (RNA Clean & Concentrator kits, ZYMO Research). Total RNA (500 ng each) was used for cDNA synthesis with the SuperScriptIII First-Strand Synthesis kit (Invitrogen). Real-time PCR was performed with the SYBR GreenER SuperMix (Invitrogen) on the QuantStudio 5 Real-Time PCR System (Thermo Fisher). Primer sequences are listed in Table S6 and their quality was assessed by the dissociation curve. Relative gene expression was determined by using the 2^-ΔΔCt^ method after normalization to the loading control GAPDH.

### In vitro kinase assay

GST or GST fusion proteins were expressed in BL21(DE3) E. coli. Cells were harvested and washed with ice-cold 1xPBS (137 mM NaCl, 10 mM Na2HPO4, 1.8 mM KH2PO4, 2.7 mM KCl, pH 7.2). Cell pellet was resuspended in ice-cold lysis buffer (1xPBS, 2 mM PMSF, 2 mM DTT, 1 x protease inhibitors (Pierce)) and sonicated (Bioruptor Pico, Diagenode) at 4LJ with alternating 10 s burst/10 s break: 10 min under high frequency and 5 min under super high frequency. Cell lysates were added with 1% Triton X-100, incubated for 15 min on ice, and cleared by centrifugation at 21,000 g for 15 min at 4LJ. Supernatant was transferred to a new tube and incubated overnight with glutathione magnetic beads at 4LJ. After washing four times with ice-cold PBS, bead-bound proteins were eluted with buffer composed of 100 mM Tris, 300 mM NaCl, 20 mM glutathione, pH 8.0. Glutathione was removed by multiple times of exchanging buffer and concentration, and the final buffer was 10 mM Tris-HCl, pH7.5. Concentration of purified proteins was measured with Nanodrop. HA-HGK or HA-HGKmt was expressed in HEK293T cells after transient transfections. Two days post transfection, cells were collected in lysis buffer (50 mM Tris-HCl, pH 7.5, 150 mM NaCl, 1% Triton X-100, 1 mM MgCl2, 1 mM PMSF, and 1 x protease inhibitor). After incubation for 15 min on ice, cell lysates were cleared by centrifugation at 21,000 g for 20 min. 20 µl anti-HA beads (Pierce PI88836) were added to the supernatants and incubated overnight at 4LJ. Beads were washed three times with 1 ml lysis buffer and twice with 1 ml kinase assay buffer (50 mM HEPES, pH7.4, 5 mM MgCl2, and 1 mM DTT). Beads in a final volume of 20 µl kinase assay buffer were supplemented with 1 µg purified GST or GST-HDAC6 proteins and 2 mM ATP. After 1 hour incubation at 37LJ, anti-HA beads were removed, and the rest of supernatant samples were separated by SDS-PAGE (10%). Protein phosphorylation was analyzed by western blotting with antibodies for phospho-Ser (pSer) or phospho-Thr (pThr).

### Pharmacokinetics

CD1 mice (21 females for Hit3 and 10 males for MAP4Ki) were injected IP with 10 mg/kg Hit3 (formulated as 5% DMSO, 5% Pharmasolve, 10% Tween 80, and 90% of 50 mM Citrate Buffer pH 4.6) or MAP4Ki (formulated in ddH2O). At the indicated timepoints after dosing, mice were sacrificed and whole blood, brain, and spinal cord were isolated. Blood samples were separated by centrifugation for 10’ at 10,000 rpm and the plasma supernatant was saved. Tissue samples were snap frozen in liquid nitrogen. Tissues were weighed and homogenized in a 4-fold volume of 1xPBS (4 x weight of brain in g = vol PBS in ml; total homogenate volume (in ml) = 5 X weight of tissue). Standards were made by spiking 50 µl blank plasma or brain homogenate with 1 µl of varying concentrations of compound and processed like samples. Brain standard curve was used to quantitate both brain and spinal cord. 50 µl of each plasma or tissue homogenate sample was crashed with 150 µl acetonitrile + 0.1% final concentration formic acid + 25 ng/ml final concentration tolbutamide IS, vortexed for 15 seconds, incubated at RT for 10 min and spun in a tabletop, chilled centrifuge for 5 minutes at 13,200 rpms. Supernatant (200 µl) was then transferred to an Eppendorf tube and spun again. Supernatant (180 µl) was analyzed by HPLC/MS.

### MAP4Ki treatments of SOD1^G93A^ mice

MAP4Ki (100 mM stock soultuion in DMSO) was diluted into 200 μM in saline. 2% DMSO in saline was prepared as vehicle. Only female mice were used for treatments since SOD1^G93A^ males frequently fought with each other when group housed. SOD1^G93A^ female mice were randomly assigned to two treatment groups (n = 12-13/group) at 40 days of age. Mice were anesthetized with isoflurane and injected intrathecally with 0.5 μl vehicle or MAP4Ki/g body weight.

### Body weight and rotarod test

Mice were weighed weekly beginning at the age of 40 days. All behavior experiments were conducted in a randomized and blinded fashion. Mice were initially trained at 38 days of age on an accelerating rotarod from Touchscreen Rota Rod (Panlab). Training was performed by placing mice on the rotarod moving at 5 rpm for 300 s. Mice were trained to stay on the rotarod for the entire 300 s. If the mouse fell from the rotarod, it was placed back until total 300 s was completed. Mice were trained for two consecutive days (89). At 40 days of age, accelerating rotarod test was conducted. The rotarod began at 4 rpm and accelerated to 40 rpm over 600 s. The time to fall was automatically recorded. Mice were tested for four consecutive trials with a resting time of 20 min between trials. Mice were examined weekly until the time at which they were unable to stay on the rotarod for more than 10 s during the four trials. Mean of the four trials was calculated as the final value.

### Survival curve

Disease progression was monitored by body weight, hind limb weakness and paralysis. The end stage was determined as the time when the mouse was unable to right itself within 15-20 s when placed on its back.

### Immunohistochemistry

Mice were sacrificed and sequentially perfused with ice-cold PBS and 4% (w/v) paraformaldehyde (PFA) in PBS. Whole spinal cords were carefully dissected out, post-fixed overnight with 4% PFA at 4°C, and cryoprotected with 30% sucrose in PBS for 48 h at 4LJ. Frozen sections of the lumbar segments were collected at 20-µm thickness by using a cryostat (Leica). Antibodies used for immunofluorescence are listed in Table S7. Nuclei were counterstained with Hoechst 33342 (Hst). CHAT^+^ neurons were analyzed in the ventral horn of the gray matter of the lumbar spinal cord segments using a Zeiss LSM 700 confocal microscope. CHAT^+^ neurons with a minimum size of 200 μm^2^ were counted as motoneurons (90). Reactive gliosis was evaluated by measuring the fluorescence intensity of GFAP or IBA1 staining in the gray matter of the spinal cords using ImageJ program. Cell counting and morphological analyses were performed in a blinded manner.

### Statistical analysis

Data are presented as mean ± SEM. Statistical significance of differences between groups was determined using a two-tailed unpaired student’s t-test or one-way ANOVA. Histological data was analyzed by one-way ANOVA and Tukey’s post hoc test. Lifespan were analyzed by Kaplan-Meier survival curve and log-rank Mantel-Cox test. Significant differences are indicated by \**p* < 0.05, \*\**p* < 0.01, \*\*\**p* < 0.001, and \*\*\*\**p* < 0.0001. All statistical tests were performed using Prism software (version 9, GraphPad).

## DECLARATIONS

### Availability of data and material

All data generated or analyzed during this study are included in the article or its supplementary information files.

### Authors’ contributions

M.-L.L., S.M., W.T., and C.-L.Z. conceived and designed the experiments. M.-L.L., S.M., W.T., X.Z., H. N., Y.Z., and J. W. performed research. C.-L.Z. supervised personnel and research. M.-L.L., S.M., W.T., and C.-L.Z. performed data analysis and interpretation. M.-L.L., S.M., and W.T. created figures for the manuscript. M.-L.L., S.M., and W.T. drafted the original manuscript. C.-L.Z revised the manuscript. All authors reviewed and approved the final manuscript.

## Acknowledgments

We thank members of the C.-L.Z. laboratory for helpful discussion, reagents and technical assistance; We also thank Dr. Andrew Lemoff at the Proteomics Core and Dr. Noelle S. Williams at the Preclinical Pharmacology Core at UTSW for technical assistance.

