## Supplemental Figures for "Chemical screens in aging-relevant human motor neurons identify MAP4Ks as therapeutic targets for amyotrophic lateral sclerosis"

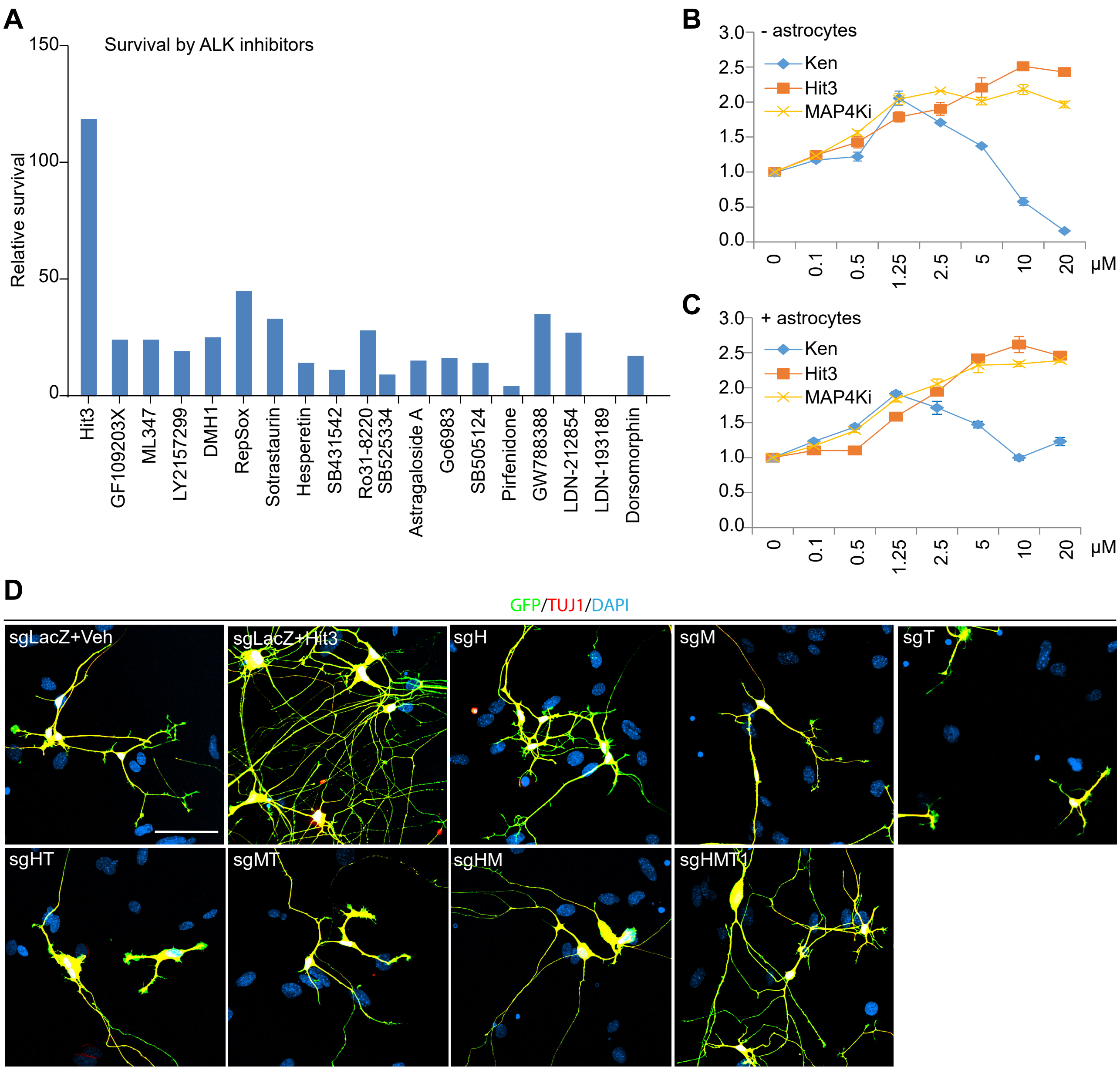


**Figure S1. MAP4K inhibition promotes survival of ALS1-hiMNs.**

A. A lack of survival effect on ALS1-hiMNs by ALK inhibitors.

B, C. Dose-dependent effect of the indicated chemicals on survival of ALS1-hiMNs with or without cocultured astrocytes (mean ± SEM; n = 4 independent samples at each concentration). Ken, kenpaullone; MAP4Ki, PF-06260933.

D. Morphological changes of ALS1-hiMNs under the indicated conditions. Genes were downregulated via sgRNAs and CRISPR-Cas9. H, HGK; M, MINK1; T, TNIK. Scale bar, 100 μm.

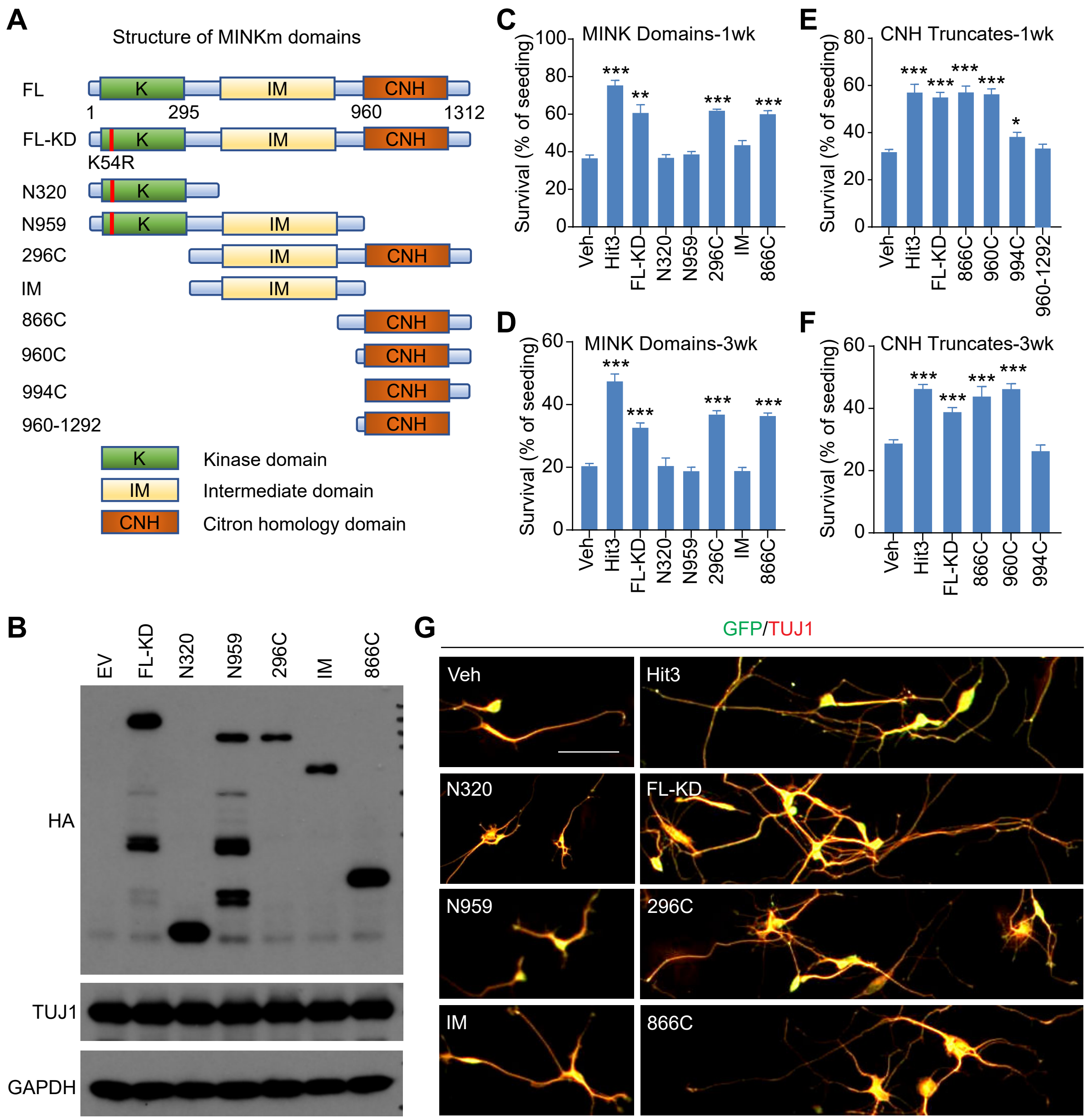


**Figure S2. CNH domain of MINK1 improves survival of ALS1-hiMNs.**

A. Schematic diagrams of MINK1 protein and its truncations. The point mutation, K54R, renders MINK1 inactive.

B. Western blotting analysis of ectopic HA-tagged proteins in ALS1-hiMNs purified at 14 dpi.

C, D. The CNH-containing domain (866C: aa866-1312) is sufficient to improve survival of ALS1-hiMNs (mean ± SEM; n = 4; **p < 0.01 and ***p < 0.001 when compared to the Veh-treated samples, Student's t-test).

E, F. The CNH domain (960C: aa960-1312) mimics Hit3's effect on survival of ALS1-hiMNs (mean ± SEM; n = 4; *p < 0.05 and ***p < 0.001 when compared to the Veh-treated samples, Student's t-test).

G. The kinase-dead MINK mutant and its functional domains recapitulated the effect of Hit3 on morphology of ALS1-hiMNs 1-week post replating. Scale bar, 100 μm.

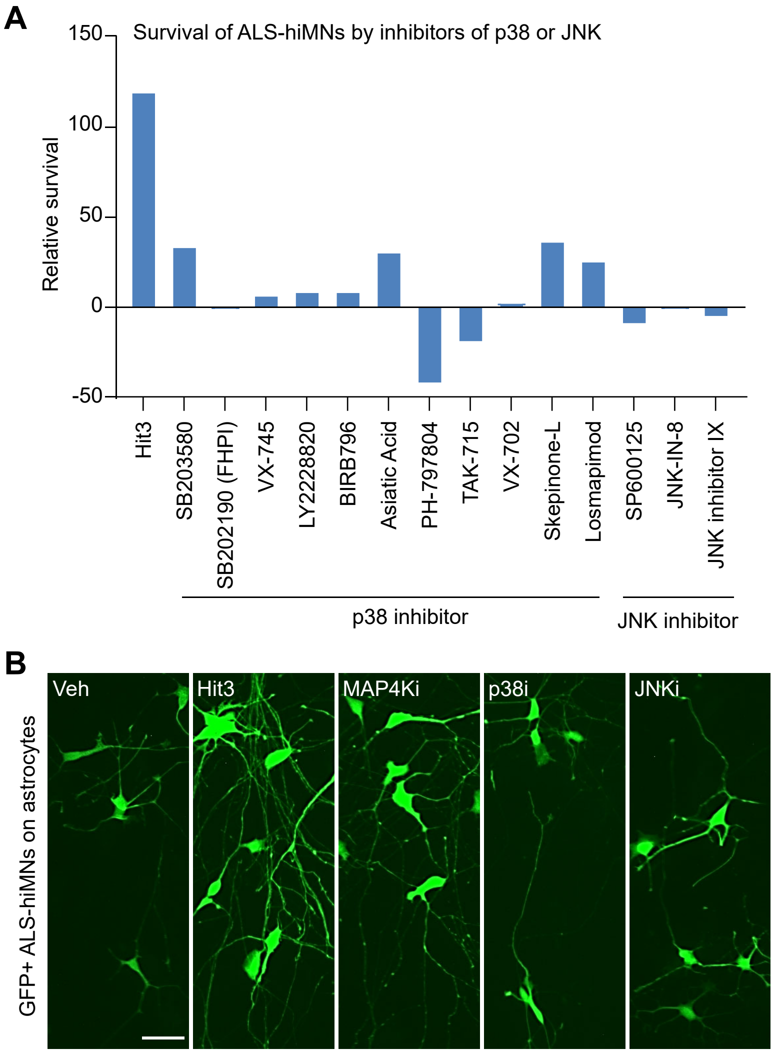


**Figure S3. A lack of protective effect of p38 or JNK inhibitors on ALS1-hiMNs.**

A. Survival assays of ALS1-hiMNs treated with the indicated inhibitors. Data were from the primary screens.

B. Effect of the selected chemicals on morphology of ALS1-hiMNs. MAP4Ki, PF-06260933; p38i, p38 inhibitor SB203580; JNKi, JNK inhibitor SP600125.

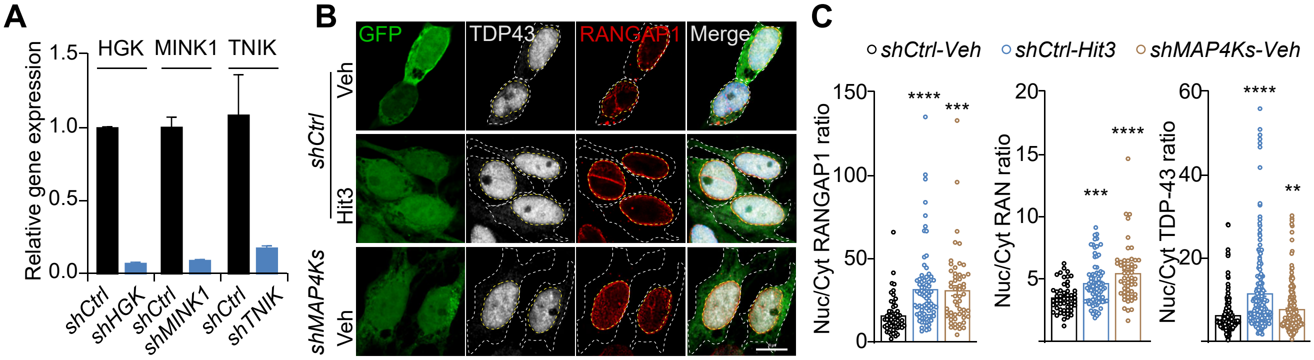


**Figure S4. Downregulation of MAP4Ks, resembling Hit3 treatments, promotes nuclear localization of key proteins in ALS1-hiMNs.**

A. qRT-PCR analysis of shRNA-mediated knockdowns in human fibroblasts.

B. Confocal images of RANGAP1 and TDP-43 in ALS1-hiMNs cocultured with astrocytes at 51 dpi. The soma and nucleus are outlined. Scale bar, 10 µm.

C. Downregulation of MAP4Ks, like Hit3 treatments, improves nuclear localization of the indicated proteins in ALS-hiMNs (mean ± SEM; $**p < 0.01$, $***p < 0.001$, and $****p < 0.0001$, one-way ANOVA).

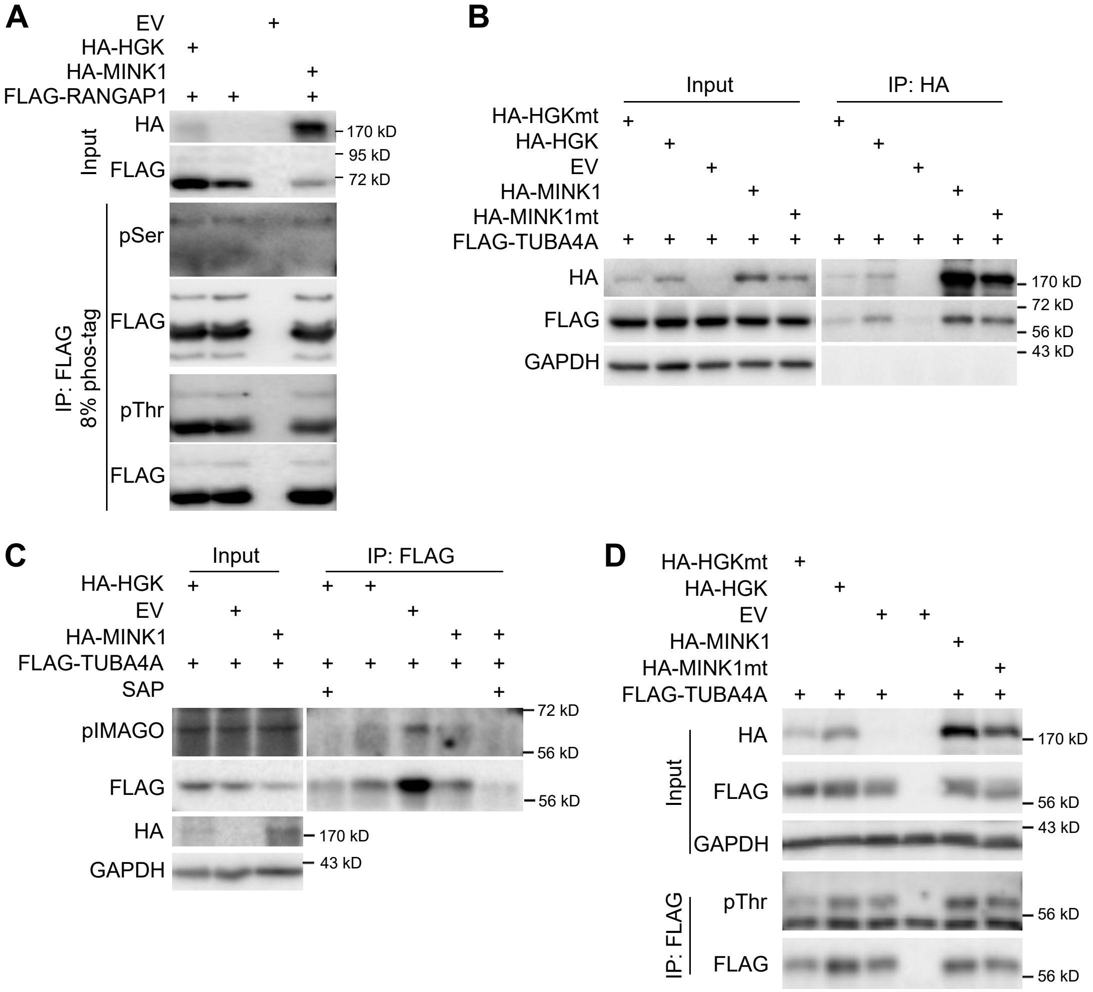


**Figure S5. Neither RANGAP1 nor TUBA4A is phosphorylated by MAP4Ks.**

A. Phos-tag SDS-PAGE and western blots failed to detect phosphorylation of RANGAP1 by MAP4Ks.

B. Co-IP results showing interactions between TUBA4A and HGK or MINK1 or their kinase-dead mutants.

C, D. pIMAGO kit or pThr phospho-antibody failed to detect phosphorylation of TUBA4A by HGK.

E, F. pIMAGO kit or pThr phospho-antibody failed to detect phosphorylation of TUBA4A by MINK1.

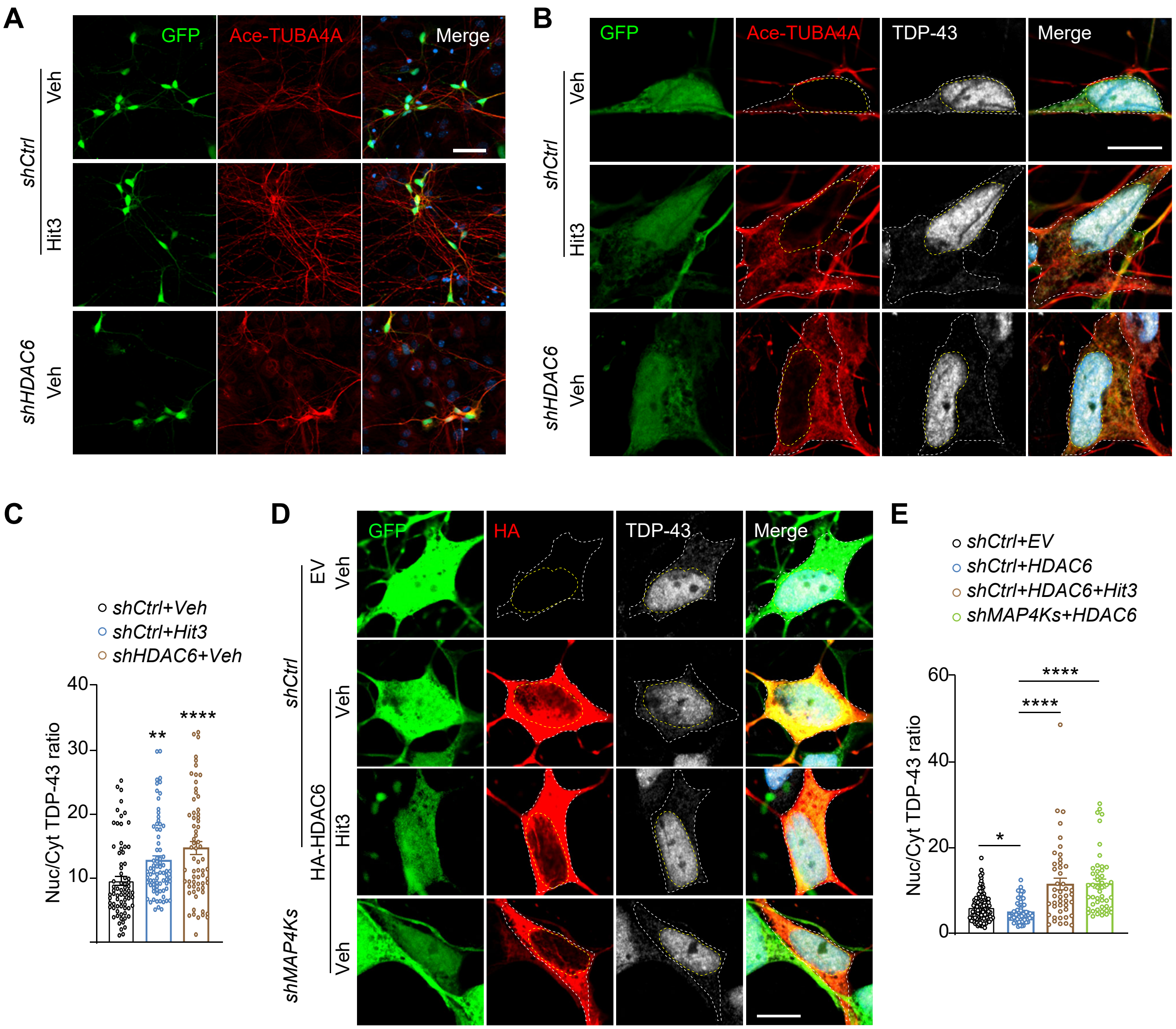


**Figure S6. HDAC6 regulates TUBA4A acetylation and TDP-43 subcellular distribution.**

A. Confocal images of ac-TUBA4A in ALS1-hiMNs cocultured with astrocytes at 28 dpi. Scale bar, 50 µm.

B. Confocal images of ac-TUBA4A and TDP-43 in ALS1-hiMNs cocultured with astrocytes at 28 dpi. Scale bar, 10 µm.

C. HDAC6 knockdown, like Hit3 treatments, improves nuclear localization of TDP-43 in ALS1-hiMNs (mean ± SEM; $**p < 0.01$ and $****p < 0.0001$, one-way ANOVA).

D. Confocal images of TDP-43 distribution in ALS1-hiMNs cocultured with astrocytes at 28 dpi. Scale bar, 10 µm.

E. Inhibition or knockdown of MAP4Ks reverses HDAC6's effect on the subcellular distribution of TDP-43 in ALS1-hiMNs (mean ± SEM; $*p < 0.05$ and $****p < 0.0001$, one-way ANOVA).

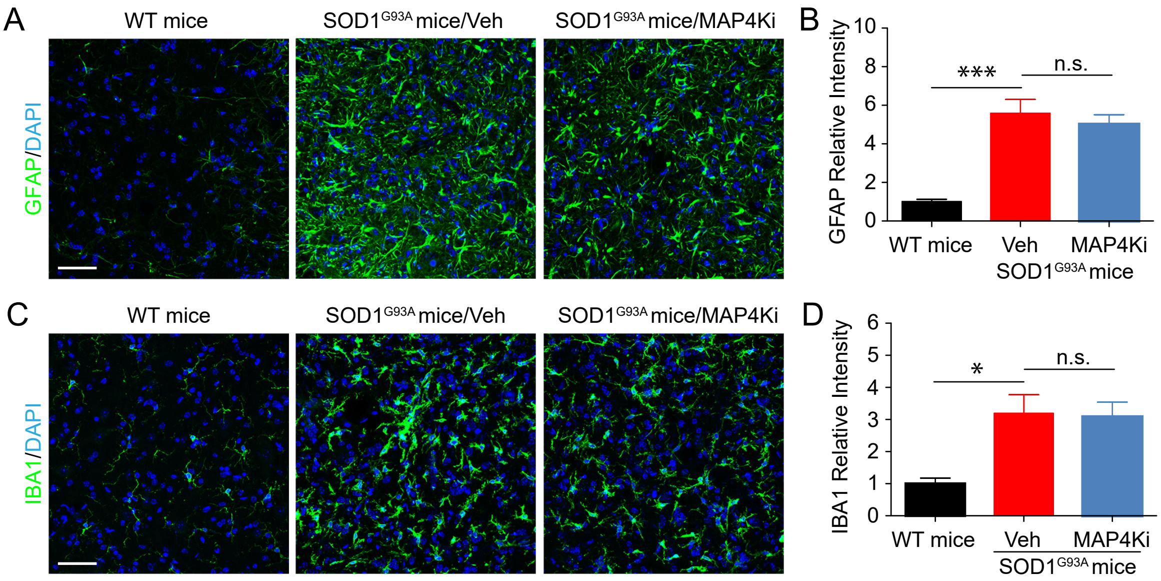


**Figure S7. MAP4Ki has no effect on reactive gliosis in SOD1$^{G93A}$ mice.**
A. Confocal images showing GFAP expression in the ventral horn of the gray matter of the lumbar spinal cord. Scale bar, 50 µm.
B. Quantification of GFAP relative intensity (mean ± SEM; n = 4-5 mice per group; ***$p < 0.001$; n.s., not significant; one-way ANOVA).
C. Confocal images showing IBA1 expression in the ventral horn of the gray matter of the lumbar spinal cord. Scale bar, 50 µm.
D. Quantification of IBA1 relative intensity (mean ± SEM; n = 4-5 mice per group; *$p < 0.05$; n.s., not significant; one-way ANOVA).
